# DNA methylation networks during pig fetal development: a joint fused ridge estimation approach

**DOI:** 10.1101/2025.05.15.654211

**Authors:** Karolina M. Wachala, Jani de Vos, Ole Madsen, Martijn F.L. Derks, Carel F.W. Peeters

## Abstract

Although an organism’s genetic information is predominantly identical among most of its cell types, the epigenome regulates the expression of the genome in a cell type- and context-dependent manner. In mammals, DNA methylation in regulatory regions, such as promoters, primarily regulates gene expression by inducing transcriptional inactivation. With genome-wide approaches came the realization that DNA methylation patterns underlying mammalian development are considerably more dynamic than previously recognized. This realization highlights the need for methodological approaches capable of capturing this phenomenon. In this study, we investigated the feasibility of modeling DNA methylation networks by jointly estimating regularized precision matrices from time- and tissue-specific omics data derived from the pig genome. For that, we utilized RNA-Seq and RRBS data that span seven pig tissues at three developmental stages: early organogenesis, late organogenesis, and newborn. Our analysis focused on 61, 48, and 74 genes — differentially expressed across developmental stages and CpG-methylated in promoter regions, from endoderm-, mesoderm-, and ectoderm-derived tissues, respectively. Using a joint fused ridge approach, we were able to *borrow* information across tissues and time points, enabling more robust network inference. This analytical framework advances exploratory methods for studying organism development using pig as a model species. Our results highlight the importance of fetal-maternal immunity and the circulatory system in early development, and shed light on dynamic interactions across tissues, organ systems, and germ layers. We anticipate that this flexible framework can be extended to other omics data and species, facilitating future research.

**Author summary:** This study explores how gene activity is controlled during pig fetal development using networks. The epigenome can turn genes on and off, depending on, e.g. the cell’s function and the organism’s growth stage. This process includes chemical changes to DNA, such as methylation. We focused on DNA methylation patterns during pig fetal development by analyzing samples from several organs at three stages of fetal growth. We combined gene activity data with DNA methylation data by using a network-based method that allows us to visualize and study how genes interact across different tissues and developmental stages. We discovered that the immune and circulatory systems are critical during early development. We have also observed complex interactions across tissues and organ systems. Our study provides a flexible toolbox with clear, step-by-step explanations. This analytical framework can be used to explore developmental patterns at the genomic scale in other species and can be adapted to different types of biological data.

## Introduction

### Significance of swine in biomedical research and livestock systems

The pig is a crucial large animal model species in various fields, such as xenotransplantation and surgical training [1]. It is used to study disease conditions owing to its anatomical and physiological similarities to humans [2]. With swine genomics recognized for its importance in biomedical research, efforts to develop complete swine (epi)genome maps have accelerated [3, 4]. Furthermore, since porcine genome sequences became crucial for discovering molecular genetic variants, they have enabled research into the genetic control of quantitative traits such as growth, reproduction, and robustness, for example, resilience to stressors, which are economically and societally desirable in livestock systems [5, 6]. The interest in genetic control of desirable traits is not surprising as pigs hold significance as a crucial global source of meat production [7]. Genomic selection, which utilizes genomic information to predict genetic potential of individual animals for desirable traits, is widely used in animal breeding with the aim of genetic improvements of livestock [8, 9]. However, as DNA methylation, among other functional epigenetic data, is pivotal to understand the regulatory mechanisms guiding pig development to e.g., predict genomic potential, efforts to characterize epigenetic mechanisms underlying porcine development are accelerating [10].

### Exploring DNA methylation and its role in swine development

Epigenetics is a bridge between an organism’s genotype and phenotype. Epigenetic modifications influence the phenotype through chemical changes altering gene expression without changing the base-pairing of the DNA sequence itself [11]. Mammals, with pigs among them, exhibit three main processes of epigenetic regulation: DNA methylation, (ii) Histone modifications, and (iii) Coding and non-coding RNAs [12]. Epigenomics relates to the analysis of these processes across numerous genes in a cell or an organism [12]. DNA methylation is a crucial epigenetic modification of gene regulation essential for mammalian development, and has been the most extensively investigated process of epigenetic regulation since its discovery in 1948 [13]. DNA methylation controls the transcriptional activity of genes by altering the chromosomal structure, DNA conformation, and DNA stability [14]. It occurs via the covalent transfer of the methyl group to the C-5 position of the cytosine ring of eukaryotic DNA, regulated by a family of DNA methyltransferases (DNMTs), namely DNMT3a and DNMT3b [15, 16]. The DNMT1 enzyme ensures the maintenance of DNA methylation. Upon establishing DNA methylation patterns, DNMT enzymes aid in continuation of said patterns into subsequent cellular generations. Although DNA methylation is usually erased during zygote formation, it is subsequently re-established in the embryo upon implementation [15]. The formation of 5-methylcytosine resulting from DNA methylation is essential for normal embryonic development in mammals, as it is vital for processes such as genomic imprinting, maintenance of X chromosome inactivation, chromosomal maintenance and genomic stability [12, 13, 15].

In the majority of mammalian somatic cells, the covalent transfer of methyl group occurs predominantly at the 5’—C—phosphate—G—3’ (CpG) dinucleotides, with most CpG sites in the genome fully methylated [17]. Additionally, gene promoters contain CpG islands, which are CpG-rich regions primarily regulating gene transcription [18]. Gene promoters are short segments of DNA (100–1,000 base pairs) where gene transcription is initiated, and methylation was revealed to regulate gene expression by switching-off transcription of the corresponding gene [19, 20]. However, the genome’s promoters remain generally unmethylated [21]. Nevertheless, owing to advancements in the genome-scale mapping of methylation, DNA methylation can be studied in various genomic contexts, from transcriptional start sites (TSSs) through gene bodies to regulatory elements, gene promoters and enhancers [19, 22]. Although traditionally DNA methylation is primarily associated with CpG islands, demethylation seems to occur predominantly in DNA regions that are not CpG islands, which has led to the recognition of DNA methylation as a vastly dynamic process [22]. The dynamic nature of DNA methylation highlights the complexity of this epigenetic regulatory process, its potential impact on gene expression, and, consequently, tissue growth and morphogenesis during organism development.

### Dynamic epigenomic switches in swine: Integrating network approaches

The GENE-SWitCH project, working in cooperation with the Functional Annotation of Animal Genomes (FAANG) Consortium, aimed to uncover the dynamic epigenetic switches in the functional genome of swine [23]. Since gene expression, transcripts and regulatory regions differ in a time- and tissue-specific manner, a static view of the genome and current genome annotation data sets may restrain a deeper understanding of gene expression and its function in mammalian development [24]. Consequently, a statistical approach allowing for the analysis of these complex and dynamic data sets is needed. However, epigenomic data are high-dimensional, as the variable dimension (genes) exceeds the observation dimension (biological replicates) [25]. The high dimensionality of the dataset leads to a so-called *curse of dimensionality* [26]. As the number of dimensions in a dataset increases, the mathematical problem space increases exponentially [27]. Therefore, the curse of dimensionality increases the computational and mathematical complexity of elucidating the epigenomics mechanisms that govern organism development [26]. In recent years, researchers developed numerous computational methods for conquering the curse of dimensionality [26]. One such method is ridge shrinkage [28]. Ridge shrinkage promotes more stable and reliable estimates of model parameters [29]. Therefore, it is applicable in omics data analyses. Next to ridge regularization, networks have gained popularity as a visually appealing and interpretable approach for the analysis of molecular mechanisms measured in thousands of variables, e.g., genes [30].

However, combining data from multiple datasets, e.g., pig tissues at developmental stages in a network, might dilute the unique biology of each sample, leading to an oversimplified representation of the dynamic epigenomics landscape during organism development. There is a need for a visually appealing and interpretable modeling approach, not only to extract DNA methylation networks and visualize how they change during organism development, but to separate signal from noise efficiently in high-dimensional DNA methylation datasets. Therefore, we aim to apply a network modeling approach utilizing data fusion over time and tissue types. Fusion will allow us to *borrow* strength from variables across DNA methylation datasets, helping to improve the stability and accuracy of generated networks, by promoting the detection of significant edges with higher precision [31]. To decide where fusion penalties should be applied, we will utilize results from hierarchical clustering of RNA sequencing (RNA-Seq) data. Moreover, using ridge shrinkage and fusion, we will extract the parameters describing network topology jointly for all tissues originating from the same somatic germ layer. Furthermore, we will visualize differential networks, capturing how DNA methylation dynamics change between developmental stages for each tissue. Thereafter, we will analyze the extracted networks. With all these steps combined our networks will more accurately capture the intricate biological context of the GENE-SWitCH Reduced Representation Bisulphite Sequencing (RRBS) datasets, allowing us to gain insights into DNA methylation dynamics driving pig development.

## Materials and methods

### Initial datasets

All the data used in this study were collected prior to our research by the

GENE-SWitCH consortium funded by the European Union’s Horizon 2020 Research and Innovation Programme. The RRBS and RNA-Seq data we used are publicly available through the FAANG data portal. RRBS uses methylation-sensitive restriction enzyme digestion on the genomic DNA [32]. The fragmented genomic DNA is then treated with bisulphite and sequenced [32]. After treatment of fragmented genomic DNA with bisulphite, unmethylated cytosine residues are converted to uracil and subsequently to thymine during polymerase chain reaction (PCR) amplification, while 5-methylcytosine residues remain unaffected [33, 34]. GENE-SWitCH has collected RRBS and RNA-Seq data from seven porcine tissues representative of the three somatic germ layers:

I. Endoderm – liver, lung, small intestine;
II. Mesoderm – kidney, skeletal muscle from hindlimb;
III. Ectoderm – hindbrain, skin.

Moreover, the data stem from three developmental stages: 30 days post-fertilization (dpf) or early organogenesis, 70dpf or late organogenesis, and from newborn piglets (NB). Additionally, at 30dpf, pooling of samples (biological replicates) coming from multiple fetuses was performed to ensure enough genetic material will be available for sequencing. RNA-Seq data collection was followed up by the normalization of read counts and differential expression analysis. The RRBS data, RNA-Seq data, and the differential expression analysis results provided by the GENE-SWitCH were the starting point for our research.

### Data preparation

The GENE-SWitCH project has utilized RRBS to study genome-wide DNA methylation of swine at a single-nucleotide resolution. Since RRBS enriches for CpG-dense regions while using restriction enzymes cutting at specific sites for fragmentation, it does not cover non-CpG areas, and CpGs in areas without the enzyme restriction site [32, 35]. Therefore, we focused on methylation in the CpG-rich promoter regions to ensure proper coverage. However, the annotation of promoter regions in CpG-methylated genes is not trivial, as for meaningful inference the most precise promoter region specification is desirable. The results from chromatin immunoprecipitation followed by sequencing (ChIP-Seq) served us to that end. This technology uses antibodies specific to a DNA-binding protein of interest to identify regulatory regions of the genome [36]. In the case of promoter regions, the H3 lysine 4 trimethylation (H3K4me3) is a core histone mark used to identify active promoter regions [36]. By investigating the distribution of histone modification signals around TSSs we identified the number of base pairs (bp) upstream and downstream of TSSs where promoters are most likely to lie. We specified that the read distribution peak is around 200bp upstream, and 50bp downstream of gene to the TSS. Moreover, we extracted methylation values only for the significantly differentially expressed genes (DEGs) between developmental stages, as obtained by GENE-SWitCH [24]. We wanted to retain all DEGs between developmental stages; therefore we did not filter on the fold change. Moreover, we excluded genes located on the X chromosome from the analyses, to mitigate potential biases related to sex differences in gene expression and methylation. We extracted the methylation values separately for each tissue at developmental stage, resulting in 21 files representing 7 pig tissues at 3 developmental stages.

Prior to exploratory analysis, we processed the data further. First, since CpG methylation might occur at multiple positions in the promoter region of a gene, while numerous genes have alternative promoters aiding pre-mRNA splicing [37], we recorded the median of replicate methylation values per gene; other data, including genes for which all the methylation values were 0, was omitted. Excluding genes that exhibited zero methylation values in all replicates ensured the presence of between-sample variation. We thereafter transposed the resulting data frames, with replicates arranged in rows, and genes arranged in columns. Secondly, we normalized (mean centered and divided by the standard deviation) the columns of the numeric matrices obtained to normalize the range of methylation values, which is a crucial step when implementing algorithms that calculate distances between data.

### Gaussian Graphical Models

We aimed to visualize networks representing relationships among variables, where each variable corresponds to the methylation value of a CpG methylated at a promoter DEG between developmental stages. Network science evolved from graph theory, a mathematical branch exploring relationships between pairs of objects [38]. Therefore, a network is just a collection of elements called nodes, linked together by their interactions, known as edges [39]. As the presence of an edge between nodes in the network indicates a conditional dependency between the variables, we aimed to extract, visualize and analyze Gaussian Graphical Models (GGMs) using the `rags2ridges` R package [40, 41]. GGM is a type of graphical model that captures pairwise relationships among variables, based on the assumption that the joint distribution of these variables follows a multivariate normal distribution [40]. When we assume multi-normality, the nodes in the network represent variables that follow a multivariate normal distribution. In this context, conditional independence between nodes is equivalent to the corresponding partial correlation zero, indicating no relationship, and therefore the absence of an edge.

Probabilistic graphs, which are networks, can depict such conditional dependencies. These graphs consist of interconnected elements, known as nodes, representing variables, while the connections between nodes, termed edges, illustrate their interactions [38]. In the case of conditional (in)dependence, nodes represent variables, whereas edges represent their conditional dependencies. If X ⊥ Y | Z Z holds true, the edge between X and Y variable will be excluded from the graph, whereas conditional dependence is illustrated by the presence of an edge, as illustrated in Fig 1.

**Fig 1.**
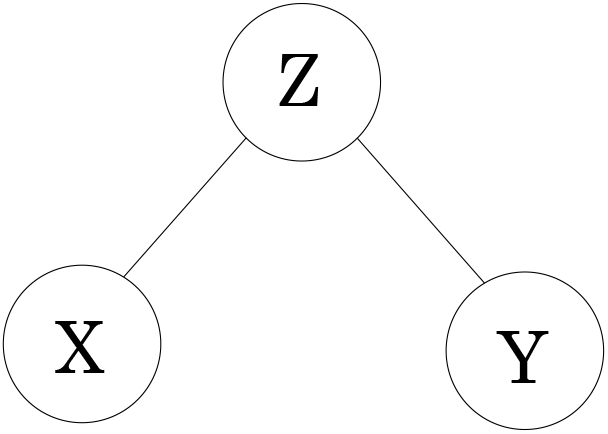
Undirected conditional independence graph. X and Y variables are conditionally independent conditioned on Z, meaning that once Z is given, knowing the value of X is superfluous concerning the value of Y, and vice versa. Since X ⊥Y |Z holds true, the edge between X and Y variable is excluded from the graph

Consequently, partial correlation can measure conditional (in)dependence [42], which can be formulated as

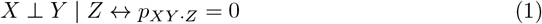

for conditional independence and

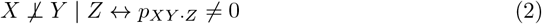

in the case of conditional dependence, where *p*_*XY ·Z*_ is a partial correlation coefficient denoting the relationship between two random variables, X and Y, conditioned on a set of random variables Z which possibly explain this association [40]. The parameters of GGMs, i.e., the partial correlations, can be estimated by inverting the sample covariance matrix Σ to obtain the precision matrix Ω. Let us consider a covariance matrix

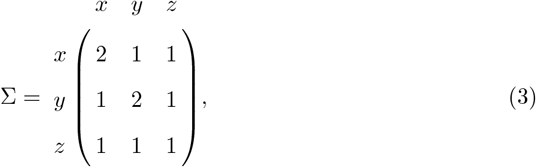

with the inverse covariance matrix (precision matrix) given by

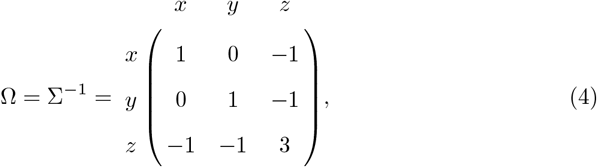

such that Σ × Ω equals the identity matrix **I**:

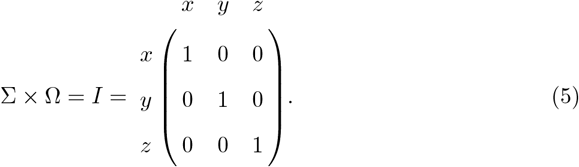

The zeros in the precision matrix indicate conditional independence of two variables – X and Y, conditioned on the third variable Z (or a set of variables). Correlation is a normalized covariance [43]

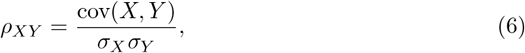

and the inversion of the covariance matrix yields a partial correlation structure between all the variables. The partial correlation coefficient 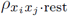 between *X*_*i*_ and *X*_*j*_ variables of a partial correlation matrix, given all other variables, is associated with the precision matrix Ω = *ω*_*ij*_ = Σ^*−*1^ such that

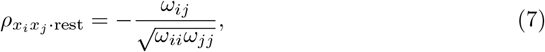

where a partial correlation matrix entry *ω*_*ij*_ = 0 indicates a zero partial correlation (absence of an edge in the network) [40].

### Nonparanormal transformation

However, GGMs assume that the modeled variables follow a multivariate normal distribution, i.e., are jointly Gaussian such that for a **X** ∈ ℝ^*n×p*^ where the rows *X*_*i*_, *i* = 1, …, *n*, are independently drawn from the same *p*-variate Gaussian distribution 𝒩 (*µ*, Σ) [44]. This assumption is not met when modeling RRBS data, since the violation of the normality assumption is frequently observed in methylation data analysis [45]. This violation originates from the fact that methylation values are bounded between 0 and 1, and their variance is usually smaller near the boundaries than near the middle of the interval, implying the violation of the homoscedasticity assumption [46]. Therefore, methylation values follow a scaled beta distribution. If not accounted for, the distribution of the project data would limit the detection of true positive edges in the networks. Consequently, gene methylation data from each tissue at developmental stage had to be transformed prior to network modeling.

The need for a multivariate Gaussian distribution between model variables implies a requirement for normality in the marginal distributions and linearity in all relationships between variables [47]. To achieve that, we transformed the RRBS data by estimating the Gaussian copula to relax the assumption of normality [48]. A copula is a specific instance of a multivariate cumulative distribution function characterized by having uniform marginal distributions, such that in a 2D (*p* = 2) setting, the marginal distributions are

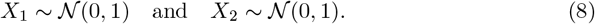

[49] proposed the nonparanormal (NPN) transformation, a special case of a Gaussian copula, to incorporate non-Gaussian data to overcome the limiting assumption of multivariate normal data. In a 2D setting, the nonparanormal transformation aims to establish a set of monotonic functions *f*_1_, *f*_2_ for a 2-dimensional random variable *X* = (*X*_1_, *X*_2_) such that the set of functions is multivariate normal distributed and *X* ∼ NPN(*µ*, Σ, *f*). This transformation preserves the conditional (in)dependence structure between the initial, non-Gaussian variables since each function *f*_*i*_ depends solely on the *X*_*i*_ variable. Therefore, the distribution of *f* (*X*) has the same factorization as the distribution of *X* [49]. With NPN transformation the uniform marginal distributions (equation 8) are transformed by a set of functions such that *f*_1_(*X*_1_) = *u*_1_ and *f*_2_(*X*_2_) = *u*_2_, with the corresponding Gaussian copula

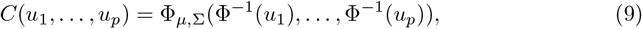

where Φ^*−*1^ denotes the inverse normal standard cumulative distribution function (CDF), and Φ_2_ denotes the CDF of multivariate distribution 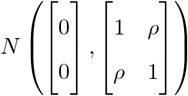 [49]. Here, we provided an explanation for a 2D setting; however, the same operation could be easily extended to higher dimensions, since the estimation of a nonparanormal transformation is computationally efficient, requiring only one pass of *X* ∈ ℝ^*n×p*^ [50]. We implemented the nonparanormal transformation using the `huge` R package, which estimates the Gaussian copula by transforming the variables marginally by applying smooth, i.e., continuously differentiable, functions [50]. We conducted the NPN transformation via the shrunken empirical cumulative distribution function (ECDF), where the parameter values are shrunken during transformation. We performed this transformation of methylation values separately for each tissue at developmental stage. This shrunken estimate helps to reduce the impact of extreme observations. Therefore, it can be effective in mitigating the impact of extreme values in situations where data values are constrained within a specific range, e.g., methylation values between 0 and 1. In this setting, we consider values approaching 0 or 1 to be extreme.

### Hierarchical clustering of RNA-Seq data

After transforming methylation values separately for each tissue at developmental stage, we performed hierarchical clustering of genes and samples using RNA-Seq data. Our goal was to visualize the correlation-based relationships between samples by utilizing gene expression data, to facilitate the construction of penalty matrices guiding joint fused ridge network extraction from methylation data. To do so, we extracted unique Ensembl gene IDs for each tissue-developmental stage combination. These genes were then matched with a TPM (transcript per million) file provided by GENE-SWitCH to extract the count-based RNA-Seq transcript expression levels of only the genes that were CpG methylated at the promoter region and differentially expressed between 30-70dpf and 70dpf-NB. Furthermore, we filtered out genes located on the X chromosome, ultimately obtaining expression values of 2718 unique genes present in all tissues at developmental stages for clustering. TPM accounts for sequencing depth, which is the total number of reads obtained from an RNA-Seq run, and gene length [51]. Nevertheless, the expression data needed pre-processing. All TPM values were log-transformed, such that *TPM* = log_2_(*TPM* + 1), to ensure that they are non-zero allowing normalization prior to clustering. Moreover, the RNA-Seq data were scaled per gene. After log transformation, the distribution of TPM-based gene expression approached a normal distribution, whereas scaling brought the variables to a comparable scale. Correlation-based hierarchical clustering followed. The pairwise similarity between genes was computed as the Pearson correlation since the expression data were approximately normally distributed. However, since scaling was performed on genes, not samples, we computed the pairwise similarity between samples as the nonparametric Spearman correlation. Both genes and samples were clustered using complete linkage to create compact, distinct clusters. Finally, we implemented the average silhouette width (AWS) method to choose the optimal number of sample clusters [52]. A silhouette represents each cluster by comparing cluster tightness and separation, illustrating which data points are positioned within the assigned cluster and which lie between the clusters [53]. AWS is calculated for each data point by measuring its intra-cluster and inter-cluster distance, with high values indicating better clustering [52]. The optimal number of clusters was chosen as k = 16, the location of the maximum, indicating tight, well-separated clusters. We subsequently generated a heatmap illustrating the hierarchical clustering of genes and samples based on the preprocessed RNA-Seq data.

### Network extraction

In preparation for network extraction, all the methylation matrices obtained after: (i) Taking a median of methylation values per gene; (ii) Removing genes with no between-sample variation; (iii) Scaling methylation values per gene; and (iv) Implementing NPN transformation, were used to obtain an intersection of genes that were present in all seven tissues and three developmental stages. However, since only one gene was shared between all the matrices, germ-layer-specific intersections of genes were obtained to allow for network modeling of methylation dynamic guiding pig fetal development. Thereafter, the methylation matrices were filtered to contain methylation values of only the genes present in the intersection. In Table 1 we provide an overview of the 21 matrices [3 tissues × 3 developmental stages + 2(2 tissues × 3 developmental stages)] obtained for further analysis.

**Table 1.**
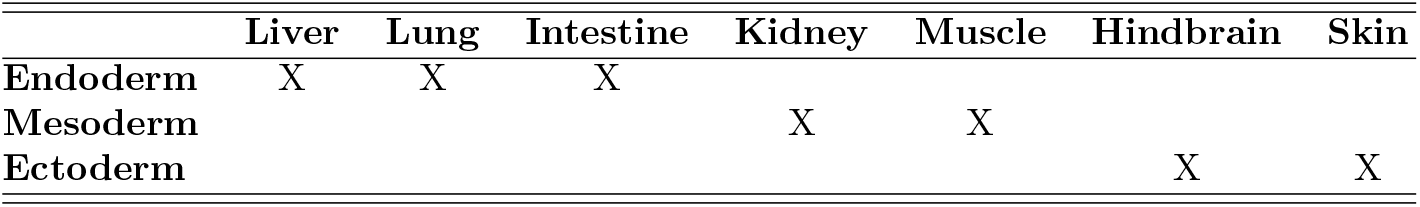
An overview of the germ layer-specific methylation matrices obtained for downstream analyses.

The methylation matrices consisted of: (i) *n* = 3, *p* = 61 for endoderm-specific matrices; *n* = 3, *p* = 48 for mesoderm-specific matrices; and (iii) *n* = 3, *p* = 74 for ectoderm-specific matrices. In Supplementary File 1, Tables S1-S3, we provide an overview of all germ layer-specific genes that we used for network modeling. We will now outline the procedure to obtain a joint estimation of germ layer- and class-specific regularized precision matrices.

First, let *y*_*ig*_ denote a realization of a *p*-dimensional random variable (methylation value of a gene), for *i* = 1, …, *n*_*g*_ independent observations (biological replicates) corresponding to *g* = 1, …, *G* non-overlapping classes (tissues at developmental stages), drawn from a multivariate normal distribution *Y*_*g*_ ∼ *N*_*p*_(0, Σ_*g*_). The parameters of the multivariate Gaussian – sample mean vector and sample precision matrix 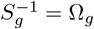 can be derived through maximum likelihood estimation (MLE). The multivariate normal log-likelihood parameterized in terms of the precision matrix for the joint 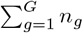 data observations is defined by

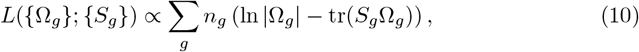

where {Ω_*g*_} and {*S*_*g*_} denote sets 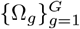 and 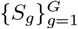 respectively, and the maximum likelihood estimate 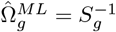 when *n*_*g*_ *> p* [41, 54]. *To derive the parameters of the* multivariate Gaussian through MLE we want to invert the sample covariance matrix *S*_*g*_ to obtain the precision matrix Ω_*g*_ for each class *g* [41]. However, it is not possible with variable dimension (*p*) exceeding the observation dimension (*n*) [41]. In such a case, the covariance matrix does not have a full rank since the!re are not enough observations to span the information of the variable dimension, and thus, it cannot be inverted. Since the variable dimension (genes) exceeds the observation dimension (biological replicates) in our data, a joint fused ridge estimation will allow the sample covariance matrix inversion, while retaining class-specific information [41]. Therefore, we seek to derive the MLE adapting formula in 10 to include the fused *l*_2_ given by

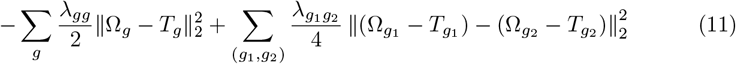

where *T*_*g*_ represent class-specific target matrices that can serve to incorporate prior information of the network structure by integrating known relationships or dependencies between variables into the matrix, *λ*_*gg*_ represent class-specific ridge penalty parameters controlling the rate of shrinkage of the unbiased, high variance Ω_*g*_ towards the biased, low variance *T*_*g*_, and 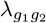 denote pair-specific fusion penalty parameters controlling the extent of entry-wise similarities between 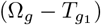 and 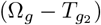 to be retained [41, 54]. Considering the fused ridge penalty in 11, the maximizing argument for a single data class *g* is defined by [54] as:

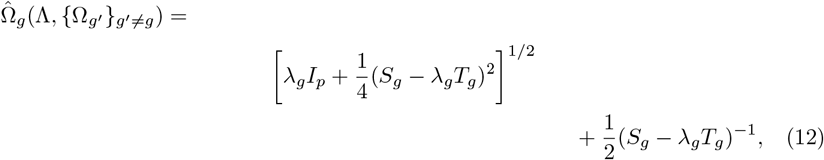

where

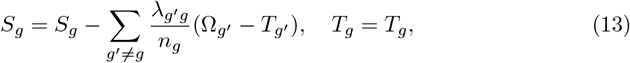

and

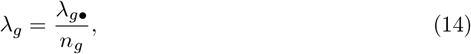

with *λ*_*g•*_ denoting the sum of entries in the *g*th row or column of the penalty matrix Λ. The penalty matrix Λ can store the class-specific (ridge) and pair-specific (fusion) penalties with *λ*_*gg*_ along the diagonal and 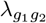 off-diagonal [41]. Since GGMs are graphs representing complex systems, e.g., methylation values of DEGs on a tissue- and time-scale, that partially share a common structure across data classes while still retaining class-specific features, setting uniform class-specific ridge penalties and uniform pair-specific fusion penalties is deemed restrictive [54, 55]. The fused *l*_2_-penalized maximum likelihood approach allows simultaneous estimation of multiple precision matrices from high-dimensional data classes. In the analysis of a factorial design, such as tissues at different developmental stages, a more precise regulation of the individual values of *λ*_*gg*_ and 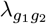 is suitable for the analysis [54]. It can be achieved by, e.g., informing the penalty matrix Λ with RNA-Seq hierarchical clustering results.

Therefore, we utilized the hierarchical clustering results to inform three separate penalty structures, such that each unique data class (tissue at developmental stage) was assigned a ridge penalty, assuming considerable differences in class-specific methylation patterns. Furthermore, fusion penalties were free to be estimated for data classes clustering the closest together. Figure providing visualization of hierarchical clustering results will follow in the Hierarchical clustering of RNA-Seq data section of the Results. Here we provide an overview of the penalty structures defined for the endoderm:

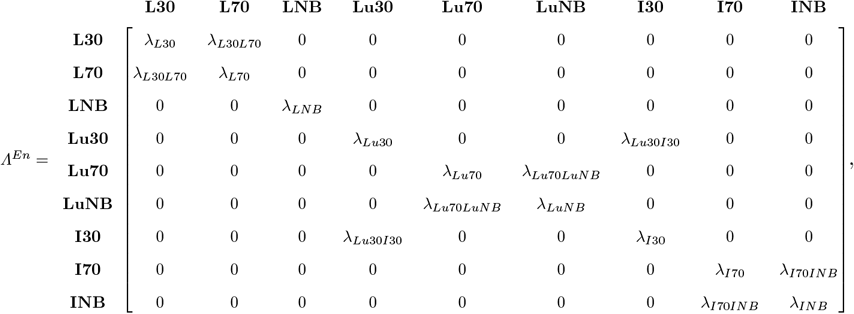

mesoderm:

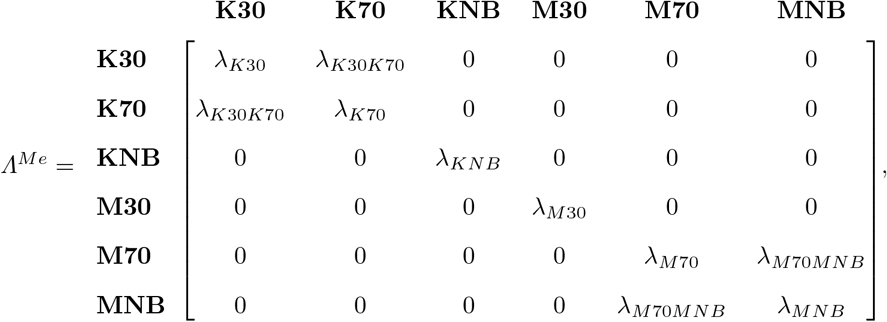

and ectoderm:

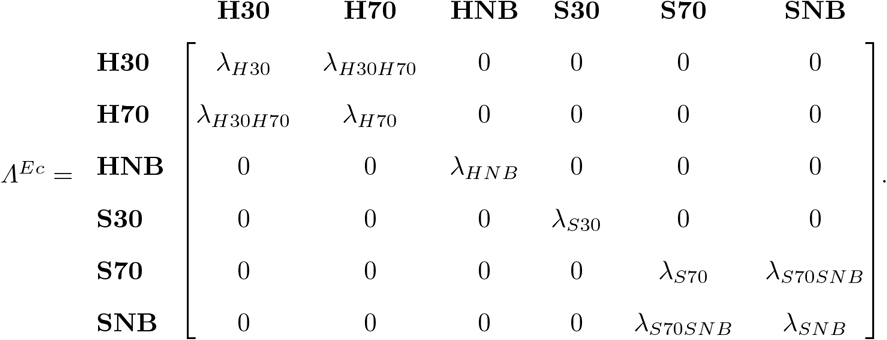

The `optPenalty.fused()` function from the `rags2ridges` R package determined each germ layer’s optimal ridge and fusion penalty parameters [41]. It utilized leave one out cross validation (LOOCV) of the negative fused log-likelihood score and employed the Nelder-Mead algorithm for function minimization [41, 56]. Thereafter, we used local false discovery rate (lFDR) with a 0.99 threshold to sparsify the regularized partial correlation matrices. The lFDR is the empirical posterior probability that an edge is null, given the observed partial correlations [41]. An edge will be retained if its posterior probability of being present is larger than or equal to the specified threshold. In our case, (1 − lFDR) ≥ 0.99. In Table 2, Table 3 and Table 4, we summarize the numbers and percentages of the edges retained for network visualization for each tissue at developmental stage.

**Table 2.** Number and percentage of edges retained for each tissue at developmental stage originating from endoderm.

|  | Liver |  |  | Lung |  |  | Intestine |  |  |
| --- | --- | --- | --- | --- | --- | --- | --- | --- | --- |
|  | 30dpf | 70dpf | NB | 30dpf | 70dpf | NB | 30dpf | 70dpf | NB |
| No. | 1271 | 1304 | 0 | 0 | 1679 | 1762 | 1292 | 0 | 1294 |
| % | 69.45 | 71.26 | 0 | 0 | 91.75 | 96.28 | 70.6 | 0 | 70.71 |

**Table 3.** Number and percentage of edges retained for each tissue at developmental stage originating from mesoderm.

|  | Muscle |  |  | Kidney |  |  |
| --- | --- | --- | --- | --- | --- | --- |
|  | 30dpf | 70dpf | NB | 30dpf | 70dpf | NB |
| No. | 0 | 0 | 744 | 756 | 741 | 0 |
| % | 0 | 0 | 65.96 | 67.02 | 65.69 | 0 |

**Table 4.**
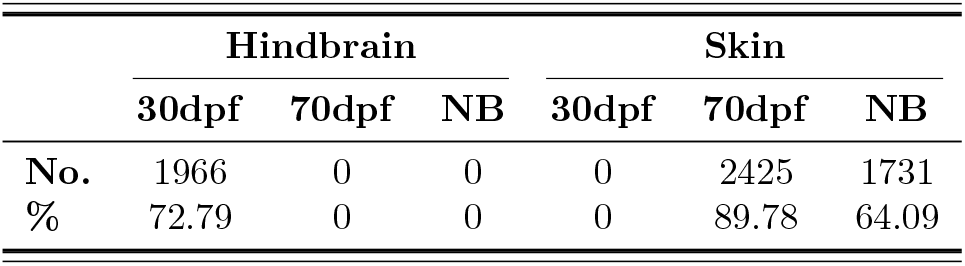
Number and percentage of edges retained for each tissue at developmental stage originating from ectoderm.

### Network visualization

Network visualization of time-dependent, tissue-specific methylation patterns followed the extraction of regularized and sparsified partial correlation matrices. We used the `Union()` function to visualize class-specific networks in the same coordinates between developmental stages for each tissue [41]. This allowed for the detection of differential connections between class-specific networks. Therefore, we obtained unions of 30-70dpf and 70dpf-NB germ-layer specific, sparsified partial correlation matrices for each tissue, sub-setting them to features containing nonzero row (column entries) so that the feature was connected in at least one of the time-specific networks. Using the `DiffGraph()` function, we visualized two differential networks (30-70dpf and 70dpf-NB) for each tissue, with the `Ugraph()` 30dpf networks determining the coordinates of 70dpf networks, whereas 70dpf networks determined the coordinates of NB networks [41]. The Fruchterman-Reingold (FR) algorithm was used to place the nodes, minimizing the number of crossing edges while ensuring that all edges were approximately equal in length [57]. To enhance the visual comparison of time-dependent differential networks, we indicated nodes (genes) to be either up-regulated (red) or down-regulated (green) between developmental stages. We extracted this information from DGE analysis results obtained by the GENE-SWitCH.

### Network analysis

Network analysis followed the visualizations of time- and tissue-dependent methylation patterns. First, we detected communities in our networks with the `cluster_fluid_communities()` function from the `igraph` R package [58, 59]. Communities in the network are nodes that are more densely connected with each other than with other nodes in the network [60]. While extracting this structural information of a network, the fluid community detection algorithm allows nodes to belong to multiple communities simultaneously [61]. Based on network topologies, we set two communities to be found for each network to capture distinct groups of nodes with cohesive interactions while maintaining graph interpretability. Second, network statistics were derived using the `GGMnetworkStats.fused()` function for each differential network [41]. In particular, we focused on genes with high betweenness centrality, indicating that these nodes are particularly important for the information flow in the networks. We did not specify a universal threshold, but rather focused on capturing a trend in the range of high betweenness centrality values in each network. We thereafter converted the gene IDs of genes with high betweennesss centrality from the Ensembl Sscrofa11.1 to Entrez IDs of human orthologs with `g:Profiler` [62]. Next, we conducted gene set enrichment analysis with `Enrichr` [63–65]. As a background, we used the single cell gene expression profiles from Developmental Single Cell Atlas of Gene RegulaTion and Expression (DESCARTES) ~ 4 million single cells from *>*110 samples representing 15 fetal organs (72–129 days post fertilization) [66].

This would allow us to evaluate if we are getting biologically feasible results for tissue-specific methylation networks obtained with the joint fused ridge approach we implemented. Thereafter, we submitted the converted gene sets with high betweennesss centrality for analysis with the ChIP-X Enrichment Analysis Version 3 (`ChEA3`) tool, to predict transcription factors (TFs) associated with the obtained methylation networks [67]. `ChEA3` computes the overlap between the submitted gene set and `ChEA3` libraries of the TF target gene set (human and mouse) with Fisher’s Exact Test, using a background of 20000 genes [67]. Benjamini-Hochberg correction is thereafter used to compute FDRs for each gene set in the library separately [67]. The correction is followed by assigning a rank to each gene set, such that 1 indicates a gene set having the lowest corrected p value from the Fisher’s Exact Test [67]. TFs are proteins regulating gene expression by binding to DNA sequence, and DNA methylation at promoters is thought to repress TFs binding [21, 68]. However, the interplay between gene expression, DNA methylation and TFs is even more complex, as it was recently proposed that TF binding can also inhibit DNA methylation [69]. Therefore, adding this additional layer of complexity to our network analysis, could provide is more complete insights into methylation dynamics driving pig fetal development.

## Results

### Hierarchical clustering of RNA-Seq data

We performed hierarchical clustering of RNA-Seq data to obtain insights into gene expression patterns underlying tissue diversification during pig fetal development. Similarities in expression profiles between samples were of interest, and we identified 16 optimal clusters with AWS (see Supplementary File 1). In Fig 2 we present the results from hierarchical clustering of the gene expression of CpG methylated at promoter regions DEGs between developmental stages, and samples representative of all seven pig tissues at three developmental stages. We excluded genes located on the X chromosome from the analysis. Moreover, we utilized the results from hierarchical clustering to decide where we should apply fusion during regularization for network extraction, therefore encoding prior, biologically meaningful information in the penalty matrices. Liver, kidney, and hindbrain samples were clustered by tissue, whereas lung, ileum, muscle, and skin clustered by developmental stage. Additionally, the following samples originating from the same germ layer clustered more closely together than with other samples: (i) Liver 30dpf – Liver 70dpf; (ii) Lung 30dpf – Ileum 30dpf; (iii) Lung 70dpf – Lung NB; (iv) Ileum 70dpf – Ileum NB; (v) Kidney 30dpf – Kidney 70dpf; (vi) Muscle 70dpf – Muscle NB; (vii) Hindbrain 30dpf – Hindbrain 70dpf; and (viii) Skin 70dpf – Skin NB. This information has been reflected in the penalty matrices, separate for each embryological layer, as shown in the Network extraction section of the Materials and methods.

**Fig 2.**
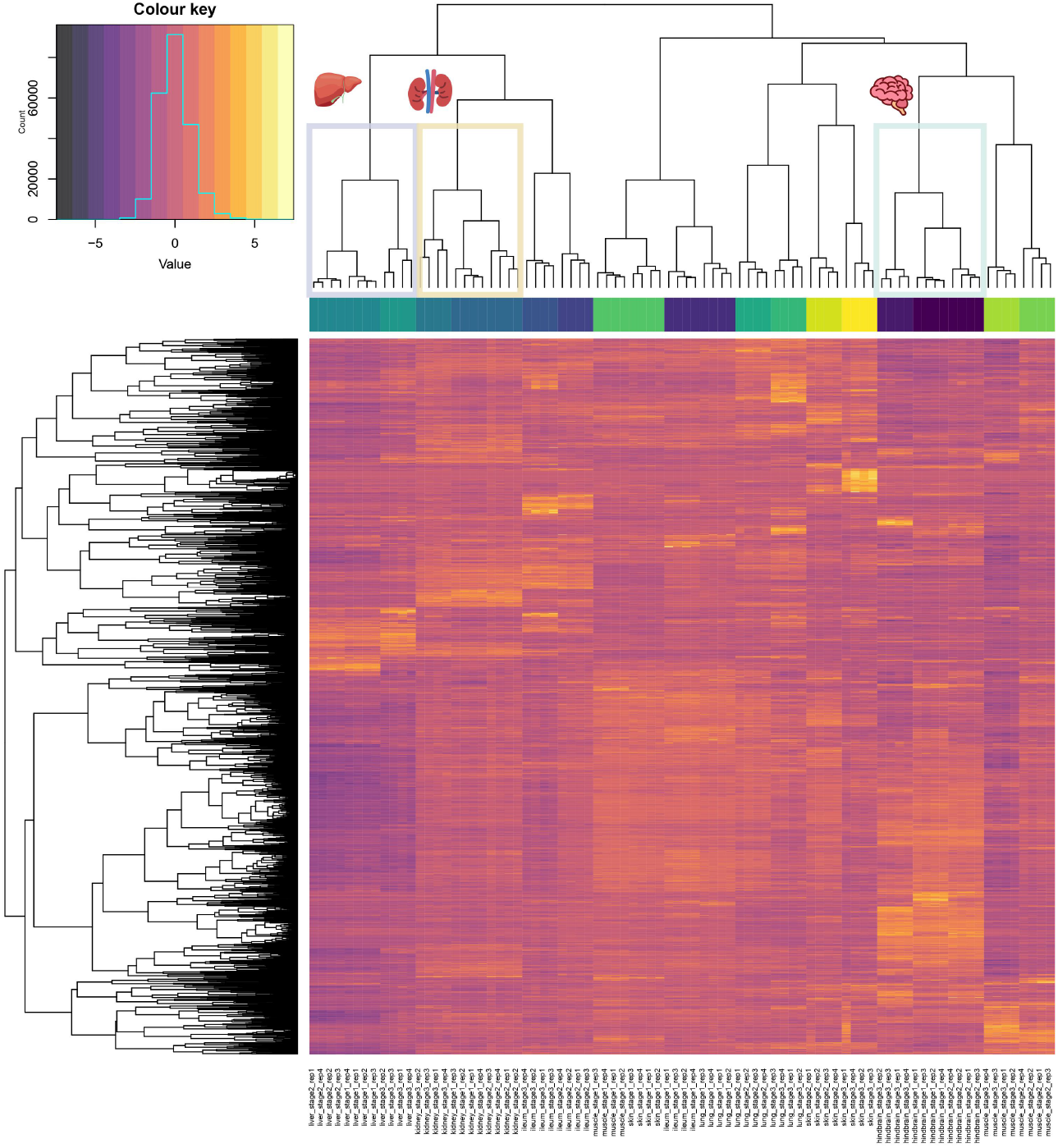
Hierarchical clustering results of TPM (Transcripts Per Million) gene expression data of all samples. Genes are clustered in rows, and samples are in columns. The row z-score corresponds to the fact that gene expression data has been mean-centered and scaled, as 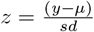 . Stage 1 denotes 30 days post-fertilization, stage 2 denotes 70 days post-fertilization, and stage 3 denotes newborn. There are four replicates for each tissue at developmental stage. Liver, kidney and hindbrain cluster by tissue, meaning that all samples coming from the same tissue cluster together

### Network extraction

Here, we present the results from jointly estimating multiple regularized partial correlation matrices under optimal penalty matrices informed by hierarchical clustering results. The optimal ridge and fusion penalty parameters estimated for each embryological layer are demonstrated. The optimal penalties were determined via LOOCV for all genes present in germ layer-specific intersections. The optimal penalties for the 61 genes found in all endoderm samples were

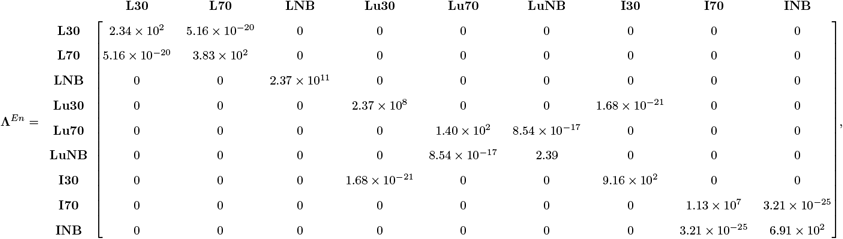

for the 48 genes present in all mesoderm samples:

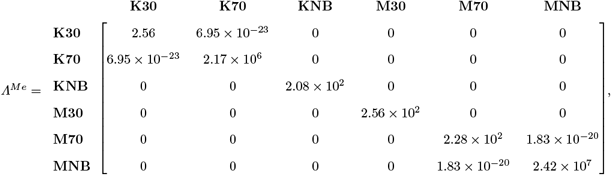

and the 74 genes present in all ectoderm samples:

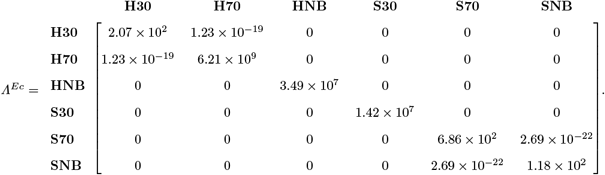

The penalty values for every germ layer-specific network extraction indicated strong differences in class-specific partial correlation matrices, favoring class-specific regularization over retaining entry-wise similarities between data classes. Moreover, we used spectral condition number plots to assess the conditioning of class-specific partial correlation matrices estimated under the optimal ridge penalties. An example plot can be found in Supplementary File 1.

### Network visualization

Following the optimal penalties extraction, conditioning assessment and support determination with a lFDR threshold of 0.99, we visualized the differential networks. The following networks were empty: Liver NB, Lung 30dpf, Intestine 70dpf, Kidney NB, Muscle 30dpf, Muscle 70dpf, Hindbrain 70dpf, Hindbrain NB, Skin 30dpf. Therefore, the differential visualizations could not be obtained for these tissues at developmental stages. Here, in Fig 3, we provide an example of a network obtained for liver between early organogenesis and late organogenesis. We present all the rest visualized differential networks in Supplementary File 1.

**Fig 3.**
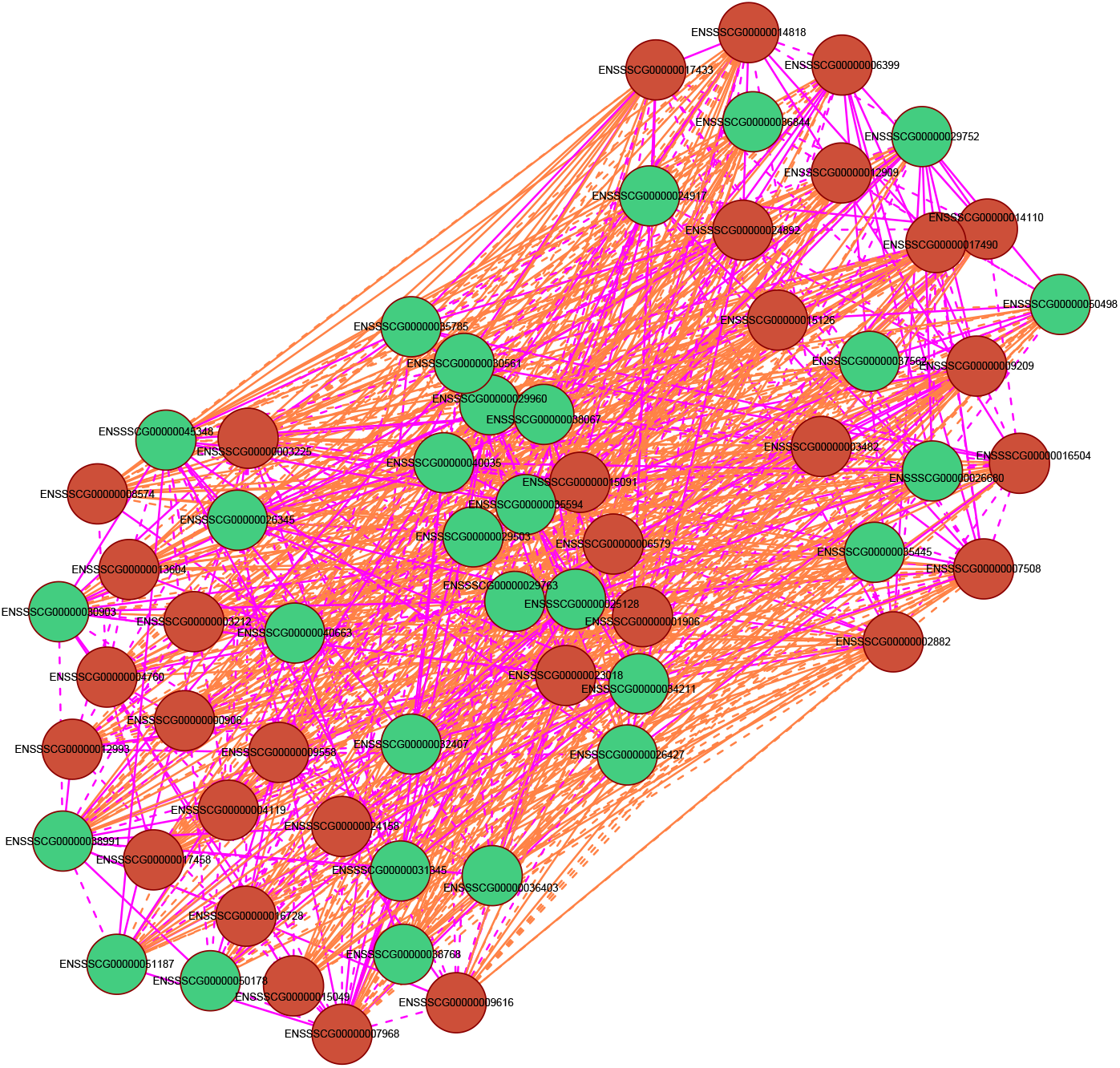
Differential network between Liver stage 30dpf and Liver stage 70dpf. Red nodes indicate up-regulated genes and green down-regulated genes. Edges unique to class Liver 30dpf are visualized in pink, while edges unique to class Liver 70dpf are in orange. Solid edges correspond to positive partial correlations, whereas dashed edges indicate negatively weighted partial correlations

The Lung 70dpf-NB and Skin 70dpf-NB networks showed a sparse *hairball* structure. Liver 30-70dpf and Skin 70dpf-NB were more densely connected than other differential networks extracted. Furthermore, there were noticeably more edges unique to late organogenesis (70dpf) in the skin network, while in the lung network edges unique to the newborn stage (NB).

### Network analysis

Continuing with network analysis, based on network topologies we decided to detect two fluid communities for each tissue developmental stage combination. In Fig 4 we provide an example of community detection visualization obtained for the Liver 30dpf network. The rest of the networks visualizing fluid community structure can be found in Supplementary File 1.

**Fig 4.**
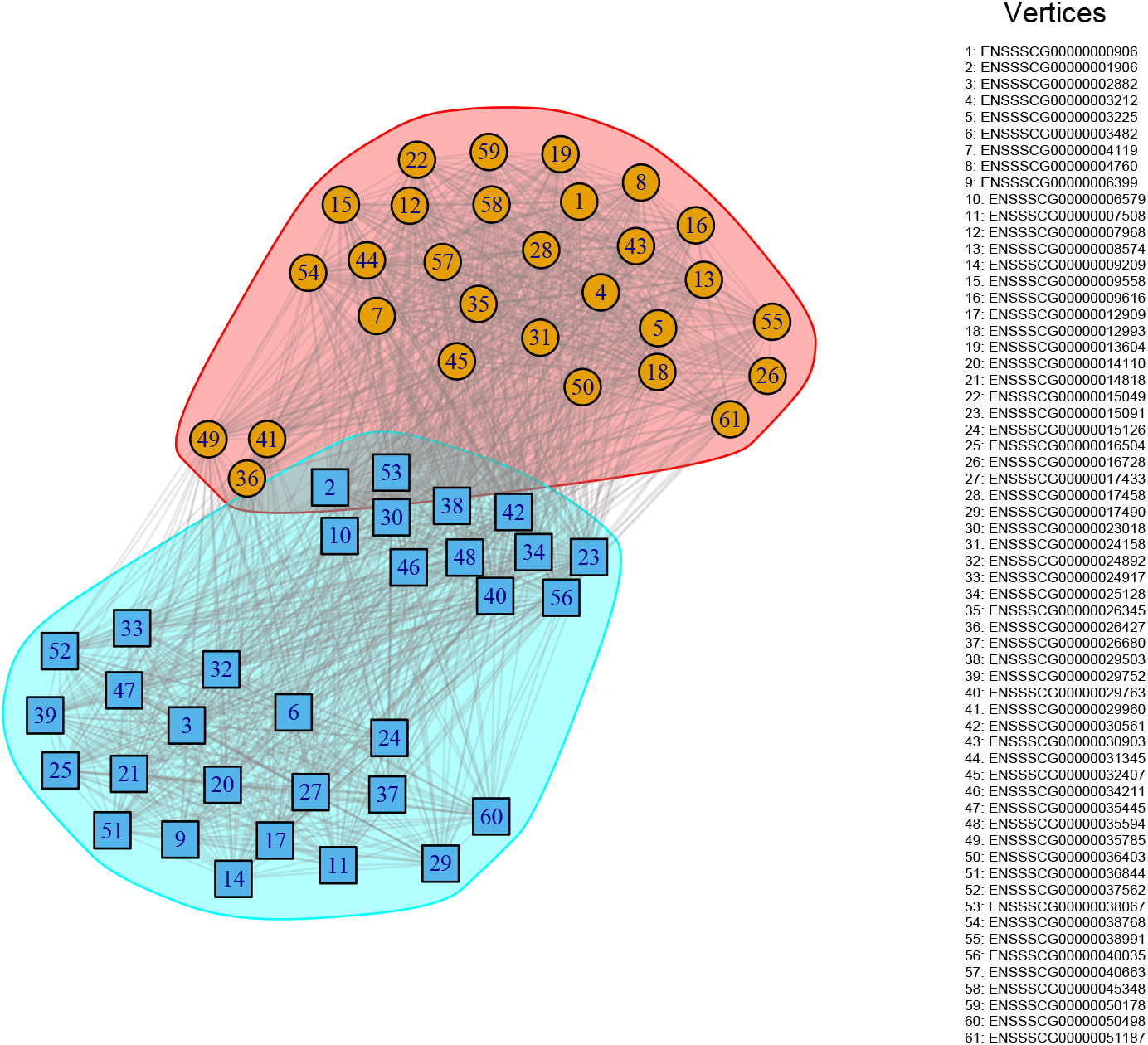
Communities detected for Liver 30dpf with the `cluster_fluid_communities()` function from the igraph R package. Each community is assigned a different colour. The legend on the right provides the encoding of the Ensembl gene IDs in the network Following community detection, we focused on analyzing network statistics. In this section, we discuss human orthologs of genes shared between the identified communities, indicating their multi-functionality. We also provide results from the gene set enrichment analyses with `Enrichr` using DESCARTES. Additionally, we present results of the TF over-representation analysis with `ChEA3`. We present our findings per tissue at developmental stage.

### Endoderm - liver

Starting with Liver 30dpf, two genes were shared between the communities, namely Cytochrome P450 family 1 subfamily A polypeptide 1 (CYP1A1) and Small Integral Membrane Protein 24 (SMIM24). While the CYP1A1 enzyme is significant throughout the entire duration of pregnancy playing a critical role in metabolizing xenobiotics within the placenta, the SMIM24 progenitor gene has been found to be expressed in the yolk sack pre-macrophages [70, 71]. The betweennesss centrality of the genes extracted for downstream analyses was between 3.45 and 33.50. Two cell types showed significant enrichment after adjusting the p-value using the Benjamini-Hochberg method, namely *Ductal Cells in Pancreas* and *Lymphoid Cells in Liver*, as seen in Table 5.

**Table 5.**
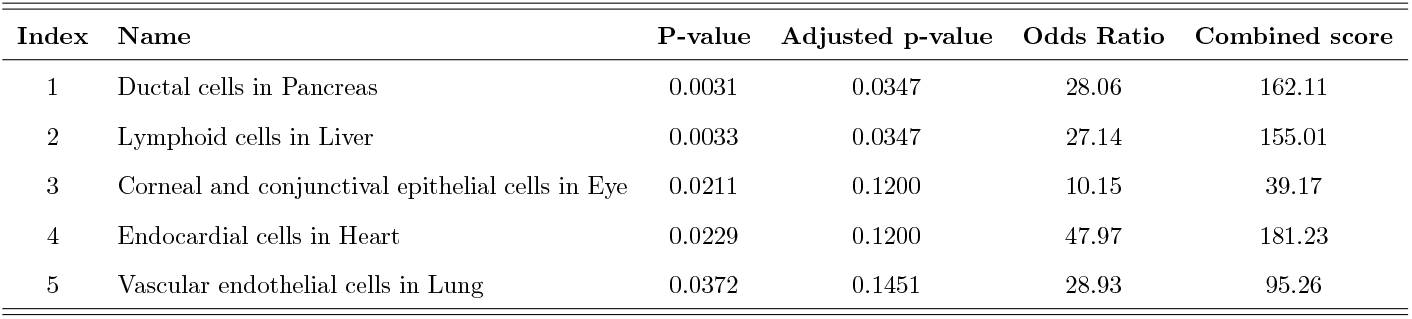
Enrichment analysis results for the Liver 30dpf network.

The enrichment for *Ductal Cells in Pancreas* is not surprising, as the endocrine, exocrine and ductal components of the pancreas originate from endoderm [72]. Furthermore, the fetal liver has been recognized as the major site of the development of the immune system, supporting the enrichment for *Lymphoid Cells in Liver* in a list of high betweennesss centrality genes from the Liver 30dpf network [73]. The innate immune lymphoid cells can also travel to other sites of organogenesis in the developing fetus through the blood [74]. Moving to the TFs, we identified KCNIP3, CREB3L3, MRFL, BARX2 and GRHL3 to be the most substantially over-represented for Liver 30dpf (see Table 6).

**Table 6.** TF target over-representation results for the Liver 30dpf network.

| Rank | TF | Mean Rank | Overlapping Genes | Gene IDs |
| --- | --- | --- | --- | --- |
| 1 | KCNIP3 | 16.50 | 2 | LMTK3, LRRC4B |
| 2 | CREB3L3 | 33.67 | 5 | F7, SMIM24, RORC, CYP1A1, CPN2 |
| 3 | MRFL | 45.00 | 2 | SMIM24, MUC2 |
| 4 | BARX2 | 50.00 | 3 | TRIM29, PRSS22, MPZL2 |
| 5 | GRHL3 | 52.33 | 3 | TRIM29, PRSS22, MPZL2 |

cAMP responsive element-binding protein 3 like 3 (CREB3L3) is known to be primarily expressed in the liver and small intestine, cooperating to maintain lipid metabolism [75]. Moreover, BarH-like homeobox 2 (BARX2) is crucial during embryonic development, being involved in the epithelial-mesenchymal interactions of organogenesis, and cytoskeletal organization [76, 77]. The expression of BARX2 is down-regulated by CpG Island hypermethylation [76].

Continuing with the next developmental stage, three genes were shared between the two fluid communities with Napsin A Aspartic Peptidase (NAPSA) among them. In the developing embryo NAPSA is expressed in mature B-cells in peripheral blood and hematopoietic stem cells in the hematopoietic bone marrow [78]. The betweenness centrality of Liver 70dpf genes extracted for downstream analyses was between 24.35 and 28.71. As presented in Table 7, the following cell types showed significant enrichment after FDR correction: *Lymphoid cells in Heart, Lymphoid cells in Lung, Lymphoid cells in Pancreas, Lymphoid cells in Intestine* and *Lymphoid cells in Adrenal*.

**Table 7.** Enrichment analysis results for the Liver 70dpf network.

| Index | Name | P-value | Adjusted p-value | Odds Ratio | Combined score |
| --- | --- | --- | --- | --- | --- |
| 1 | Lymphoid cells in Heart | 0.0039 | 0.04758 | 24.54 | 136.34 |
| 2 | Lymphoid cells in Lung | 0.0071 | 0.0476 | 17.87 | 88.52 |
| 3 | Lymphoid cells in Pancreas | 0.0072 | 0.0476 | 17.63 | 86.88 |
| 4 | Lymphoid cells in Intestine | 0.0076 | 0.0476 | 17.17 | 83.74 |
| 5 | Lymphoid cells in Adrenal | 0.0097 | 0.0485 | 15.09 | 69.95 |

Although the adrenal medulla originates from mesoderm, while the adrenal cortex from ectoderm, we already established that innate immune lymphoid cells can travel between organogenesis sites [73, 79, 80]. Moreover, the fetal liver has been shown to accommodate the progenitors of various types of lymphoid cells [73]. Next, we identified IRPF9, ZNF831, BATF2, SCML4 and TBX21 to be the most substantially over-represented TFs for Liver 70dpf (see Table 8). Interferon regulatory factor 9 (IRF9) belongs to the IRF TF family involved in the transcriptional regulation of the immune system and cell growth, working at the intersection of innate and adaptive immunity [81]. Moreover, Basic leucine zipper ATF-like transcription factor 2 (BATF2) has been found to be a key factor in myeloid differentiation, therefore regulating immune response [82].

**Table 8.** TF target over-representation results for the Liver 70dpf network.

| Rank | TF | Mean Rank | Overlapping Genes | Gene IDs |
| --- | --- | --- | --- | --- |
| 1 | IRF9 | 14.75 | 7 | NAPSA, IFI35, PTPRCAP, GPR37L1, MYO1F, HERPUD1 |
| 2 | ZNF831 | 16.33 | 3 | GPR171, PTPRCAP, MYO1F |
| 3 | BATF2 | 27.33 | 3 | NAPSA, IFI35, MYO1F |
| 4 | SCML4 | 28.33 | 3 | GPR171, PTPRCAP, MYO1F |
| 5 | TBX21 | 29.25 | 5 | NAPSA, GPR171, PTPRCAP, MYO1F, HERPUD1 |

### Endoderm - lung

Moving on to the lung networks, we identified five genes to be shared between the fluid communities. Two of them are particularly interesting, namely Coagulation Factor X (F10) and Thromboxane A Synthase 1 (TBXAS1). Both are involved in hemostasis, with F10 encoding the vitamin K-dependent coagulation factor X [83, 84]. TBXAS1 is a member of the cytochrome P450 superfamily of enzymes, similar to CYP1A1 introduced in the Liver 30dpf section [84]. Since both lung and liver originate from the endoderm, this is in line with the hematopoietic function of the liver during fetal development. The betweenness centrality of the top genes from the Lung 70dpf network was between 3.26 and 3.51. We identified no significantly enriched cell types after adjusting the p-value using the Benjamini-Hochberg method. However, without the correction (see Table 9), similar to Liver 30dpf, in Lung 70dpf network *Ductal cells in Pancreas* was the most enriched term.

**Table 9.** Enrichment analysis results for the Lung 70dpf network.

| Index | Name | P-value | Adjusted p-value | Odds Ratio | Combined score |
| --- | --- | --- | --- | --- | --- |
| 1 | Ductal cells in Pancreas | 0.0057 | 0.1428 | 19.81 | 102.29 |
| 2 | PAEP MECOM positive cells in Placenta | 0.0123 | 0.1541 | 13.16 | 57.87 |
| 3 | Vascular endothelial cells in Lung | 0.0501 | 0.2805 | 20.89 | 62.54 |
| 4 | Squamous epithelial cells in Stomach | 0.0663 | 0.2805 | 15.58 | 42.28 |
| 5 | Lymphoid cells in Heart | 0.0995 | 0.2805 | 10.13 | 23.37 |

The second most significant term was *PAEP MECOM positive cells in Placenta* (p-value = 0.01232). In male human fetus both *PAEP* -positive and *MECOM* -positive cells in the placenta is associated with the expression of the *Xist* or *Tsix* gene [66]. *Xist* encodes a long non-coding RNA (lncRNA) that coats one of the X chromosomes in females, leading X chromosome inactivation to achieve dosage compensation between genders [85]. *Tsix* is a lncRNA antisense to *Xist* [86]. The presence of these markers is of maternal origin, and while *Xist* corresponds to maternal decidualized stromal cells, *Tsix* corresponds to maternal endometrial epithelial cells [66]. Next, we identified FOXN1, RBPJL, NR1D1, TBPL2 and CENPB to be the most substantially ovverepresented TFs for Lung 70dpf (see Table 10). Forkhead box protein N1 (FOXN1) is responsible for thymus organogenesis, by regulating thymic epithelial cells (TECs), therefore contributing to in utero T-cell development, and aiding the immune response [87–89].

**Table 10.**
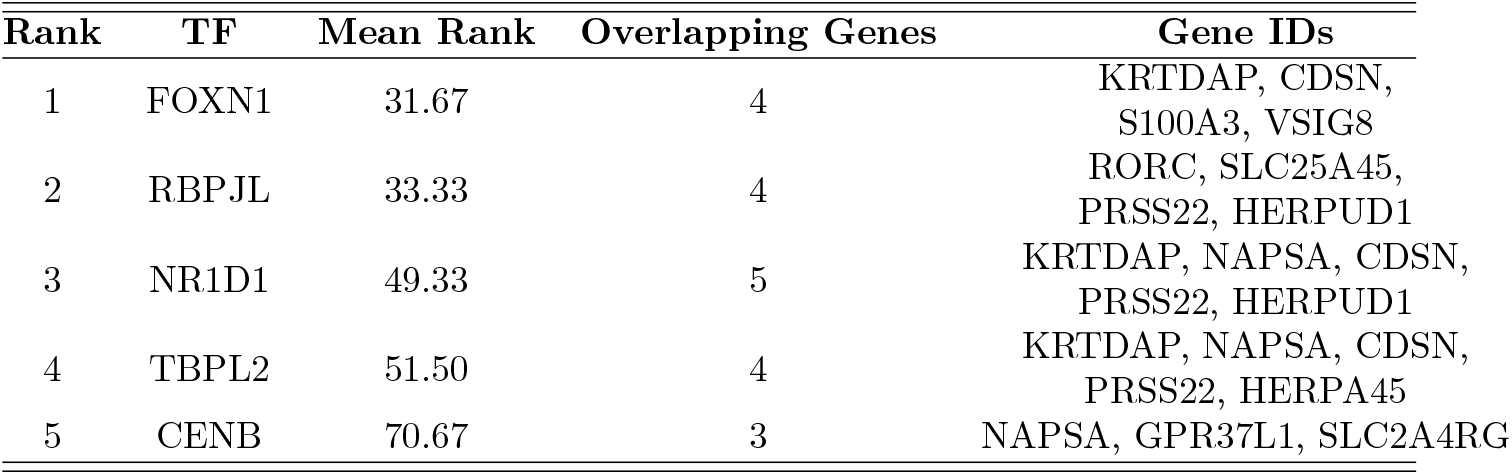
TF target over-representation results for the Lung 70dpf network.

Next, the Lung NB network community detection resulted in the most overlapping nodes (26 out of 61 present) from all networks. Therefore, we decided to conduct gene ontology enrichment analysis of the human orthologs of these 26 genes using `g:Profiler` [62]. However, we did not obtain any significant results after FDR correction. Nevertheless, one should keep in mind that this is a relatively small gene set. Continuing with nodes with the highest betweenness centrality, the same 26 genes had betweenness centrality equal to 1.67, the lowest from all the tissues at developmental stages. After FDR correction, no terms showed significant enrichment. The cell types with the highest p value were *Ductal cells in Pancreas* (0.01056), similar to Liver 30dpf and Lung 70dpf, and *IGFBP1 DKK1 positive cells in Placenta* (0.01107), as shown in Table 11.

**Table 11.** Enrichment analysis results for the Lung NB network.

| Index | Name | P-value | Adjusted p-value | Odds Ratio | Combined score |
| --- | --- | --- | --- | --- | --- |
| 1 | Ductal cells in Pancreas | 0.0106 | 0.1273 | 14.02 | 63.81 |
| 2 | IGFBP1 DKK1 positive cells in Placenta | 0.0111 | 0.1273 | 13.67 | 61.57 |
| 3 | AFP ALB positive cells in Spleen | 0.0418 | 0.2450 | 6.60 | 20.95 |
| 4 | PDE1C ACSM3 positive cells in Stomach | 0.0470 | 0.2450 | 22.15 | 67.72 |
| 5 | Vascular endothelial cells in Lung | 0.0679 | 0.2450 | 15.03 | 40.43 |

Insulin-like growth factor-binding protein 1 (IGFBP1), and Dickkopf-related protein 1 (DKK1) are marker genes associated with decidual cells, which the decidua (pregnant endometrium) consists of, while the immune cells are one of the main cell types making up the decidual cells [90, 91]. We subsequently identified OSR1, NR1I3, BATF2, IRF9, and IRF7 to be the most overrepresented TFs for Lung NB (see Table 12).

**Table 12.** TF target over-representation results for the Lung NB network.

| Rank | TF | Mean Rank | Overlapping Genes | Gene IDs |
| --- | --- | --- | --- | --- |
| 1 | OSR1 | 26.00 | 4 | RHBDF1, ANO1, FAM180A |
| 2 | NR1I3 | 29.33 | 4 | DMGDH, IGFBP1, F7, CYP1A1, SLC25A45 |
| 3 | BATF2 | 34.00 | 5 | ZBP1, NAPSA, IFI35 |
| 4 | IRF9 | 50.00 | 4 | ZBP1, NAPSA, IFI35, GPR37L1, SLC25A45 |
| 5 | IRF7 | 54.33 | 3 | ZBP1, IFI35 |

Odd skipped-related 1 (OSR1) is associated with connective tissue formation and myogenesis in limbs, pointing towards mesoderm development, whereas *constitutive androstane receptor* (NR1I3) expression is enriched in adult mammalian liver and its expression in human embryonic stem cells has been linked to differentiation and maturation of hepatic-like cells [92, 93].

### Mesoderm - kidney

Continuing with mesoderm networks, we identified six genes shared between the fluid communities for Kidney 30dpf. On the basis of our previously presented results, we decided to focus on two of them, namely Cytoglobin (CYGB) and the previously introduced CYP1A1. CYGB is unique to vertebrates, encoding a protein called hexacoordinate hemoglobin, distinguishing it from pentacoordinate hemoglobins primarily engaged in oxygen transportation [94]. With its binding mechanism for ligands varying significantly from pentacoordinate hemoglobins, the CYGB protein is believed to safeguard cells during periods of oxidative stress [94]. CYGB has also been revealed to activate hepatic stellate cells (HSC) during disturbances in oxygen homeostasis, aiding in liver fibrogenesis [95]. Furthermore, CYGB has been shown to protect kidney fibroblasts against oxidative stress caused by a lack of blood supply; and therefore, a lack of oxygen [96]. As for CYP1A1, its expression in the kidney increases in response to oxidative stress, whereas in the liver it is involved in the oxidation of fatty acids and toxins [97, 98]. Next, we looked at 11 genes with the highest betweenness centrality (between 27.82 and 32.32) in the Kidney 30dpf network. We did not find any statistically significant terms after FDR correction. The top enriched cell types after correction were *Smooth Muscle Cells in Heart, Oligodendrocytes in Cerebellum, Vascular endothelial cells in Placenta, Lymphoid cells in Lung*, and *Lymphoid cells in Placenta* (Table 13).

**Table 13.** Enrichment analysis results for the Kidney 30dpf network.

| Index | Name | P-value | Adjusted p-value | Odds Ratio | Combined score |
| --- | --- | --- | --- | --- | --- |
| 1 | Smooth muscle cells in Heart | 0.0198 | 0.0887 | 56.84 | 222.86 |
| 2 | Oligodendrocytes in Cerebellum | 0.0484 | 0.0887 | 22.55 | 68.29 |
| 3 | Vascular endothelial cells in Placenta | 0.0513 | 0.0887 | 21.25 | 63.11 |
| 4 | Lymphoid cells in Lung | 0.0725 | 0.0887 | 14.80 | 38.82 |
| 5 | Lymphoid cells in Placenta | 0.0739 | 0.0887 | 14.50 | 37.77 |

The enrichment for *Smooth Muscle Cells in Heart* is not surprising, since smooth muscle cells (SMCs) play a significant role in the structural and functional makeup of numerous organs throughout embryonic development, while the heart is of a mesodermal origin [99, 100]. Moreover, oglidendrocytes wrap myelin, allowing fast and energy efficient neuronal signaling, and nerve-muscle communication governs voluntary movements and breathing [101, 102]. Like the kidney and heart, muscles originate from the mesoderm, while cerebellum and the nervous system from the ectoderm [99, 103]. The development of the nervous system from the ectoderm is initiated by a signal from mesoderm [103]. Additionally, vascular endothelial cells (EC) form a single layer lining the interior of arteries, veins, and capillaries, facilitating development and maintenance of the circulatory system. The endothelium functions as an endocrine, i.e., hormonal organ as well [104, 105]. Coupling this information with the observed enrichment for lymphoid-associated cell types supports our results pointing towards the importance of the immune system and interactions between various tissues (and germ layers) during fetal development. Eventually, we investigated the top enriched TFs for genes with high betweenness centrality in the Kidney 30dpf network. In Table 14, we present the results from this enrichment analysis.

**Table 14.**
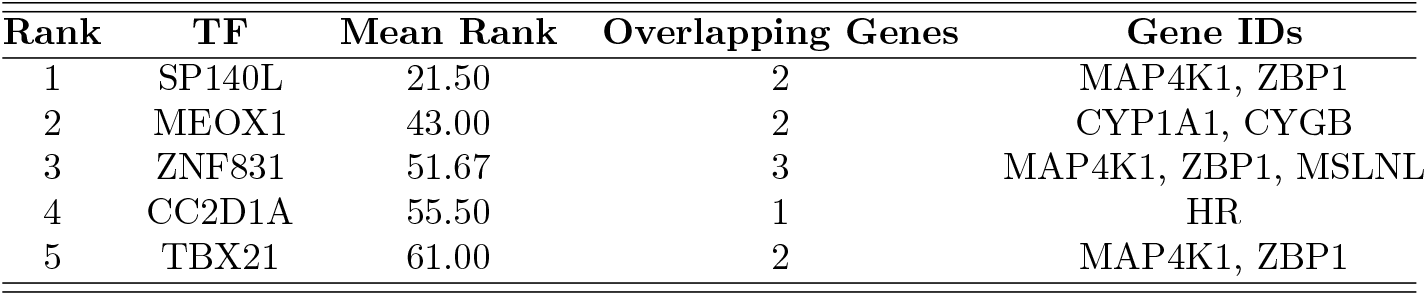
TF target over-representation results for the Kidney 30dpf network.

The top enriched TFs for Kidney 30dpf were SP140L, MEOX1, ZNF831, CC2D1A, and TBX21. The speckled protein (SP) family members are expressed primarily in leukocytes, and the SP140 locus has been associated with multiple autoimmune diseases, such as multiple sclerosis (MS) damaging the central nervous system, which is considered to be a T-cell-mediated disease [106]. Furthermore, mesenchyme homeobox 1 (MEOX1) has been identified as a positive regulator of smooth muscle cell differentiation during embryogenesis in mice [107].

With respect to the Kidney 70dpf genes, we did not identify any genes that were shared between the two fluid communities. The betweenness centrality of 10 genes extracted for downstream analyses was between 26.60 and 38.48. After applying FDR correction, two cell types showed a significant enrichment, namely *Cardiomyocytes in Heart* and *Megakaryocytes in Muscle* (see Table 15).

**Table 15.** Enrichment analysis results for the Kidney 70dpf network.

| Index | Name | P-value | Adjusted p-value | Odds Ratio | Combined score |
| --- | --- | --- | --- | --- | --- |
| 1 | Cardiomyocytes in Heart | 0.007973 | 0.03446 | 147.96 | 714.91 |
| 2 | Megakaryocytes in Muscle | 0.008616 | 0.03446 | 17.22 | 81.88 |
| 3 | Retinal pigment cells in Eye | 0.04603 | 0.1003 | 23.77 | 73.18 |
| 4 | Chromaffin cells in Adrenal | 0.05129 | 0.1003 | 21.25 | 63.11 |
| 5 | SATB2 LRRC7 positive cells in Heart | 0.06267 | 0.1003 | 17.24 | 47.76 |

Cardiomyocytes are cardiac muscle cells responsible for involuntary contractions of the heart, supplying the organism with blood [108]. Megakaryotes are rare cells found in the bone marrow, responsible for the production of platelets, which aid the formation of blood clots and wound healing [109]. In addition to assisting thrombosis, platelets are know to influence the immune response as well by, e.g., directing T-cell activation and differentiation [110, 111]. The circulatory, renal and musculoskeletal systems originate from the mesoderm, and our results point towards the importance of interactions between different tissues during fetal development [99]. Focusing on TF over-representation analysis, the top five overrepresented TFs for Kidney 70dpf were BORCS8-MEF2B, SNAI3, NKX2-5, HLF and THRA (see Table 16).

**Table 16.** TF target over-representation results for the Kidney 70dpf network.

| Rank | TF | Mean Rank | Overlapping Genes | Gene IDs |
| --- | --- | --- | --- | --- |
| 1 | BORCS8-MEF2B | 51.00 | 2 | RGS14, UNC13D |
| 2 | SNAI3 | 53.67 | 3 | RGS14, TRIB3, UNC13D |
| 3 | NKX2-5 | 58.00 | 1 | RNF207 |
| 4 | HLF | 66.67 | 1 | TEF |
| 5 | THRA | 74.75 | 2 | TEF, LRRTM2 |

Snail family transcriptional repressor 3 (SNAI3) is a member of the SNAIL gene family, known to be involved in orchestrating mesoderm formation during embryogenesis [112], whereas NK2 homeobox 5 (NKX2-5) encodes a transcription factor involved in heart formation [113].

### Ectoderm - skin

Continuing with the Skin 70dpf networks, we identified two genes: solute carrier family 6 member 7 (SLC6A7) and lymphocyte specific protein 1 (LSP1) to be shared between the computed fluid communities. SLC6A7 belongs to the gamma-aminobutyric acid neurotransmitter gene family [114]. It is also responsible for the production of L-proline transporter protein in the mammalian brain [114]. The presence of this gene in the skin network is not surprising, since both the skin and the brain originate from the ectoderm [115]. Additionally, LSP1 encodes a protein expressed in macrophages, lymphocytes, and the endothelium; therefore, it is involved in the immune response [116]. Moving to the 16 genes with the highest betweenness centrality in the Skin 70dpf network (values between 6.40 and 6.52), we did not obtain any significant cell type enrichment results (see Table 17). However, one must keep in mind that we continuously worked with small gene sets in this study, therefore it is not surprising.

**Table 17.** Enrichment analysis results for the Skin 70dpf network.

| Index | Name | P-value | Adjusted p-value | Odds Ratio | Combined score |
| --- | --- | --- | --- | --- | --- |
| 1 | Squamous epithelial cells in Stomach | 0.0560 | 0.2812 | 18.70 | 53.86 |
| 2 | Megakaryocytes in Adrenal | 0.0830 | 0.2812 | 12.38 | 30.82 |
| 3 | Megakaryocytes in Heart | 0.0889 | 0.2812 | 11.52 | 27.88 |
| 4 | Ductal cells in Pancreas | 0.0918 | 0.2812 | 11.13 | 26.57 |
| 5 | Smooth muscle cells in Eye | 0.1027 | 0.2812 | 9.88 | 22.47 |

The top five overrepresented TFs we identified for Skin 70dpf genes were KCNIP3, EBF4, BATF2, SOX9, and ARID5A (Table 18).

**Table 18.**
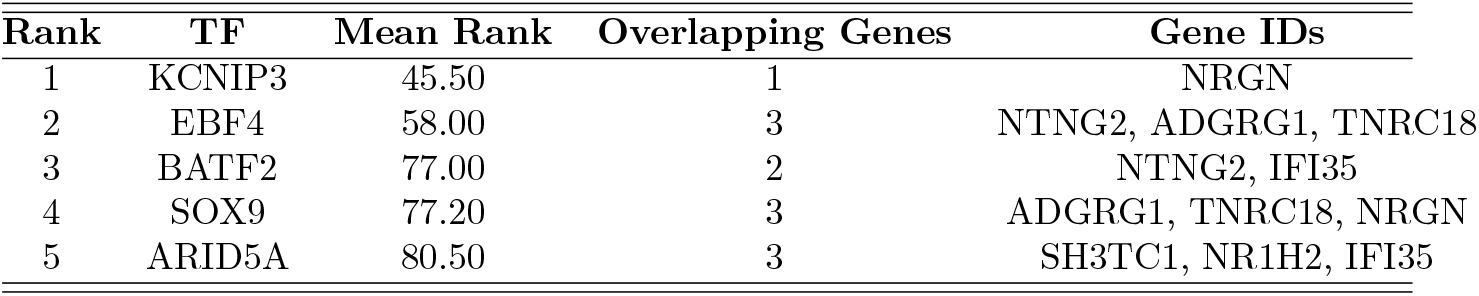
TF target over-representation results for the Skin 70dpf network.

We have already introduced BATF2 in the Liver 70dpf section as a regulator of the immune response. Furthermore, we found that Early B cell factor 4 (EBF4) is expressed in human immune cells, having a cytotoxic lymphocyte function, therefore aiding the immune response [117]. Additionally, SRY-box transcription factor 9 (SOX9) TF is known to be involved in a variety of developmental processes. Not only does SOX9 expression determine cell fate, driving tissue diversification in all germ layers, but it is also a crucial regulator of sex determination during mammalian embryonic development [118, 119].

Finally, we focused on analyzing the Skin NB network. Seven genes were found at the intersection of the two extracted communities. Here, we introduce fucose mutarotase (FUOM): an inhibitor of neuron differentiation [120] that is up-regulated between 70dpf and NB in our differential network visualization (see Supplementary File 1 Fig. S6, here as ENSSSCG00000010791). Moreover, we looked at 9 genes with the highest betweenness centrality in the Skin NB network. Their values were high compared to other tissues at developmental stages (between 62 and 74.2), however, we identified only one significantly enriched cell type after FDR correction, namely *Megakaryocytes in Muscle* (Table 19). We discussed megakaryotes in the Kidney 70dpf section.

**Table 19.** Enrichment analysis results for the Skin NB network.

| Index | Name | P-value | Adjusted p-value | Odds Ratio | Combined score |
| --- | --- | --- | --- | --- | --- |
| 1 | Megakaryocytes in Muscle | 0.0069 | 0.0348 | 19.69 | 97.79 |
| 2 | Skeletal muscle cells in Eye | 0.0919 | 0.1714 | 11.66 | 27.84 |
| 3 | AFP ALB positive cells in Spleen | 0.1075 | 0.1714 | 9.87 | 22.02 |
| 4 | Corneal and conjunctival epithelial cells in Eye | 0.1371 | 0.1714 | 7.59 | 15.08 |
| 5 | Hepatoblasts in Liver | 0.1923 | 0.1923 | 5.21 | 8.60 |

Thereafter, we looked at overrepresented TFs in Skin NB networks, and identified ZNF605, ZNF154, ZNF221, ZNF780B, and ZNF626 (Table 20).

**Table 20.** TF target over-representation results for the Skin NB network.

| Rank | TF | Mean Rank | Overlapping Genes | Gene IDs |
| --- | --- | --- | --- | --- |
| 1 | ZNF605 | 19.33 | 3 | ZNF37A, PEG10, PCBD2 |
| 2 | ZNF154 | 66.67 | 3 | ZNF37A, GPX7, PCBD2 |
| 3 | ZNF221 | 72.00 | 1 | ZNF37A |
| 4 | ZNF780B | 93.00 | 1 | ZNF37A |
| 5 | ZNF626 | 98.67 | 3 | ZNF37A, PEG10, PCBD2 |

Interestingly, all the top five TFs belong to the zinc finder proteins (ZNF) family, which is the largest TF family in the human genome [121]. In humans ZNF605, ZNF154, ZNF780B, and ZNF626 exhibit the highest expression in the thyroid (with the thyroid hormones modulating a variety of immune cells), whereas ZNF221 in the brain (originating from ectoderm, similar to skin) [115, 122, 123].

## Discussion

As the omics revolution picks up pace, the need for novel approaches to analyzing high-throughput biological data ever-increasing in complexity and size, which can conquer the curse of dimensionality, becomes evident. At the same time, model interpretability and visually appealing analytical approaches are desirable. Therefore, we aimed to determine whether the joint fused ridge network extraction informed by the results from hierarchical clustering of RNA-Seq data can ensure the extraction of the relevant data signal in network modeling of DNA methylation data coming from multiple swine tissues and developmental stages.

The findings concerning hierarchical clustering of the RNA-Seq data revealed that liver, kidney and hindbrain samples clustered by tissue, whereas lung, ileum, muscle and skin clustered by developmental stage. This aligns with the results obtained by the GENE-SWitCH, even though we worked with a subset of gene expression data, that is, CpG methylated at promoter regions DEGs, excluding genes located on the X chromosome [24]. Additionally, at 30dpf we observed the lung and ileum, which originate from endoderm, clustering together, suggesting that during early organogenesis, they have not yet fully differentiated.

Subsequently, we informed the penalty structures with results from hierarchical clustering to estimate the optimal penalty matrices for each germ layer separately. Ideally, we would have specified one common penalty structure for all tissues and developmental stages together; however, only one gene was shared among all data classes. In that sense, our analysis might be considered reductionist since we have ignored interplay between tissues originating from different germ layers. We observed favoring of class-specific regularization (ridge) over retainment of entry-wise similarities between data classes (fusion) in all germ layer-specific optimal penalty matrices. This suggests that all data classes (tissues at developmental stages) are quite distinct concerning methylation patterns, further emphasizing the dynamic nature of the (epi)genome. The highest class-specific ridge penalty was evaluated by LOOCV for Liver NB, indicating that the precision matrix approached the target matrix, suggesting negligible signal in the data. We also observed high ridge penalties for Lung 30dpf, Intestine 70dpf, Muscle 70dpf, Hindbrain 70dpf, Hindbrain NB, and Skin 30dpf, explaining empty networks obtained for the above-mentioned tissues at developmental stages. However, Kidney 70dpf also had a relatively high ridge penalty compared to other data classes (*λ*_*gg*_ = 2.17 × 10^6^), but we were able to extract the network. Notably, all partial correlation matrices estimated under the optimal penalty parameters determined for germ layer-specific network extraction were well-conditioned, as suggested by condition number plots. This illustrates that the *l*_2_ regularization procedure implemented with `rags2ridges` led to well-conditioned estimates of the partial correlation matrices for network visualization and downstream analyses.

Continuing with cell type enrichment and TF over-representation analysis, using gene sets with the highest betweenness centrality from each data class specific network, we observed biologically feasible results for each tissue. Concerning the recurring trend of immune-response associated terms for all tissues, a fetus is considered vulnerable to infections in utero [124]. However, the maternal-fetal immunological landscape involving, e.g., the placenta, lymphoid cells, and megakaryotes, supports the developing fetus in all stages of pregnancy [125]. The fact that immune cells in mammals are vertically transmitted from the mother to the fetus in utero, would also explain the observed enrichment for terms associated with the placenta [126]. This transmission promotes neonatal immunity, aiding the newborn in resistance against infections [126]. Additionally, a recurring trend of enrichment for terms associated with the circulatory system, further supports our observations, since various types of immune cells are found in blood, circulating through the developing organism [127]. This would also explain why we obtain enrichment for a different tissue that we are analyzing at times, as every tissue belongs to an organ system dependent on nutrient and oxygen supply by the blood for healthy fetal development [128]. The lung and liver are the exceptions, not being sustained by the fetal blood workflow; however, their waste products and carbon dioxide will enter the fetal circulatory system [128]. The results obtained suggests that our analytical framework allows for extraction of biologically feasible data signal from high-dimensional methylation data spanning seven swine tissues at three developmental stages.

However, our results should be approached with caution due to the limitations of contemporary omics research and the methodological constraints of the project itself. The sample size is the major issue to consider when aiming to accurately estimate network parameters and, therefore, the network topology. As introduced before, omics data analyses often suffer from the *curse of dimensionality*. Consequently, we implemented joint fused ridge estimation. Nevertheless, conquering singularity does not solve the issues of difficulty in collecting pig tissue samples, especially at early developmental stages. The considerations for ethical animal experimentation call for the adoption of the 4Rs principles (Reduction, Refinement, Replacement and Responsibility) [129], which in essence, require the least feasible number of biological replicates to be collected. Moreover, there is often not enough genetic material in a single sample (which resulted in pooling of samples at 30dpf), and the desired sample itself is hard to extract. To that end, a developing field of organoids, which are lab-grown cell-based in vitro models mimicking the functioning of in-vivo tissues, arises as a valuable alternative to traditional animal experiments [130]. Furthermore, genes involved in early development are often less well annotated than genes expressed in mature tissues. Additionally, literature on swine development at genomics resolution is scarce; therefore we conducted cell type enrichment and TF over-representation analyses on the human orthologs. Coupled with the small size of the gene sets analyzed, the insights obtained from our networks might not represent early developmental stages in swine accurately, since we reviewed literature focusing on human and mice development. These limitations highlight the importance and novelty of the efforts of projects such as GENE-SWitCH. We believe that our analyses would benefit from utilizing (epi)genomics data measured at single-cell resolution, therefore accounting for tissue heterogenity, allowing us to extract networks that more accurately represent methylation dynamics during pig fetal development.

Nevertheless, one should realize that in this study, we did not test any biological hypothesis, but rather, we propose a novel exploratory analysis approach to extract a mechanistic view of the genes and their interactions based on methylation values. For that reason, we developed an interpretable and visually appealing workflow for the analysis of omics datasets spanning various tissues and developmental stages. And although there were limitations associated with gene annotations at early developmental stages, literature availability and the dataset itself, we were able to extract biologically feasible results with our approach.

Future research would likely benefit from utilizing single-cell omics data to account for tissue heterogeneity. In that setting, we would recommend fusion between cell populations within tissues, not the tissues themselves. Additionally, to minimize biological noise in the collected samples, organoids could be considered. However, the question of how closely can organoids mimic the functioning of in-vivo tissues at a molecular level, remains unanswered. Moreover, single-cell sequencing and organoids, however promising, are an expensive alternative to more traditional technologies. With respect to the joint fused ridge, we identified a lack of fusion penalty parameter assessment methods available, which could open avenues for future research.

Nevertheless, our proposed methodology could be implemented in other omics fields. The fused ridge approach has been used successfully in multiple omics domains, such as metabolomics, to study blood-based metabolomic signatures in Alzheimer’s disease, and proteomics to identify sepsis biomarkers from patients’ serum [131, 132]. We also see potential in exploring differences between tissues (differential networks between tissues), instead of focusing on differential molecular dynamics between developmental stages.

## Conclusion

To conclude, we demonstrated that:

I. Utilizing RNA-Seq hierarchical clustering results to control the individual values of the ridge and fusion penalties is a feasible approach to incorporate biologically meaningful information in the penalty structures of time- and tissue-dependent swine DNA methylation networks;
II. Joint fused ridge regularization procedure with the use of `rags2ridges` R package allows for the extraction of the relevant data signal in an ill-posed problem, namely network modeling of DNA methylation data subjected to class membership.

Furthermore, the results from our network analysis point toward the importance of the immune and circulatory system in the developing fetus, shedding light on the dynamic *network* of interactions between tissues, organ systems and germ layers.

## List of Abbreviations

DNMT: DNA methyltransferase
CpG: 5’—C—phosphate—G—3’
TSS: Transcription Start Site
FAANG: Functional Annotation of Animal Genomes consortium
RNA-Seq: RNA Sequencing
RRBS: Reduced Representation Bisulphite Sequencing
PCR: Polymerase Chain Reaction
30dpf: 30 days post fertilization
70dpf: 70 days post fertilization
NB: Newborn
ChIP-Seq: Chromatin Immunoprecipitation assays with Sequncing
H3K4me3: Histone 3 Lysine 4 trimethylation
bp: Base pair
DEG: Differentially Expressed Gene
GGM: Gaussian Graphical Model
NPN: Nonparanormal
CDF: Culmulative Distribution Function
ECDF: Empirical Culmulative Distribution Function
TPM: Transcripts Per Million
AWS: Average Silhouette Width
MLE: Maximum Likelihood Estimation
LOOCV: Leave One Out Cross Validation
lFDR: Local False Discovery Rate
FR algorithm: Fruchterman-Reingold algorithm
DESCARTES: Developmental Single Cell Atlas of Gene RegulaTion and Expression
TF: Transcription Factor
ChEA3: ChIP-X Enrichment Analysis Version 3
CYP1A1: Cytochrome P450 family 1 subfamily A polypeptide 1
SMIM24: and Small Integral Membrane Protein 24
CREB3L3: cAMP responsive element-binding protein 3 like 3
BARX2: BarH-like homeobox 2
NAPSA: Napsin A Aspartic Peptidase
IRF9: Interferon regulatory factor 9
BATF2: Basic leucine zipper ATF-like transcription factor 2
F10: Coagulation Factor XThromboxane A Synthase 1
TBXAS1: Thromboxane A synthase 1
lncRNA: Long non-coding RNA
FOXN1: Forkhead box protein N1
IGFBP1: Insulin-like growth factor-binding protein 1
DKK1: Dickkopf-related protein 1
OSR1: Odd skipped-related 1
NR1I3: Constitutive androstane receptor
CYGB: Cytoglobin
SMC: Smooth muscle cell
EC: Endothelial cells
SP: Speckled protein
MEOX1: Mesenchyme homeobox 1
SNAI3: Snail family transcriptional repressor 3
NKX2-5: NK2 homeobox 5
SLC6A7: Solute carrier family 6 member 7
LSP1: Lymphocyte specific protein 1
EBF4: Early B cell factor 4
SOX9: SRY-box transcription factor 9
FUOM: Fucose mutarotase
ZNF: Zinc finder protein
4Rs: Reduction, Refinement, Replacement, Responsibility

## Supporting information

**S1 File**. Supplementary File 1 contains additional graphs and tables as indicated in manuscript text.

## Statements and Declarations

### Ethics approval and consent to participate

Not applicable

### Consent for publication

Not applicable

### Availability of data and materials

The data used in this study were collected by the GENE-SWitCH

(https://www.gene-switch.eu/) and are available through the FAANG data portal at https://data.faang.org/projects/GENE-SWitCH. The accession code is: (i) PRJEB41822 for RRBS; (ii) PRJEB41970 for RNA-Seq; and (iii) PREJEB70458 for ChIP-Seq. The results supporting the conclusions of this article are included in Supplementary File 1. The code and generated, pre-processed data are available through a GitHub repository at https://github.com/wachalak/methylation-networks/.

### Author’s contributions

KMW contributed to the study’s design, processed and analyzed the data, interpreted the results, and drafted the manuscript. JdV supervised KMW, contributed to data generation, and initial data analysis. OM conceived the GENE-SWitCH methylation study and supervised JdV. MFLD supervised KMW, assisted with data analysis and manuscript editing. CFWP supervised KMW, contributed to the study’s conceptualization, supported results interpretation and manuscript editing. All authors reviewed and approved the final manuscript.

## Acknowledgments

Not applicable

