## Supplementary File 1 for "DNA methylation networks during pig fetal development: a joint fused ridge estimation approach"

**† Shared last authorship.**

### FIGURES

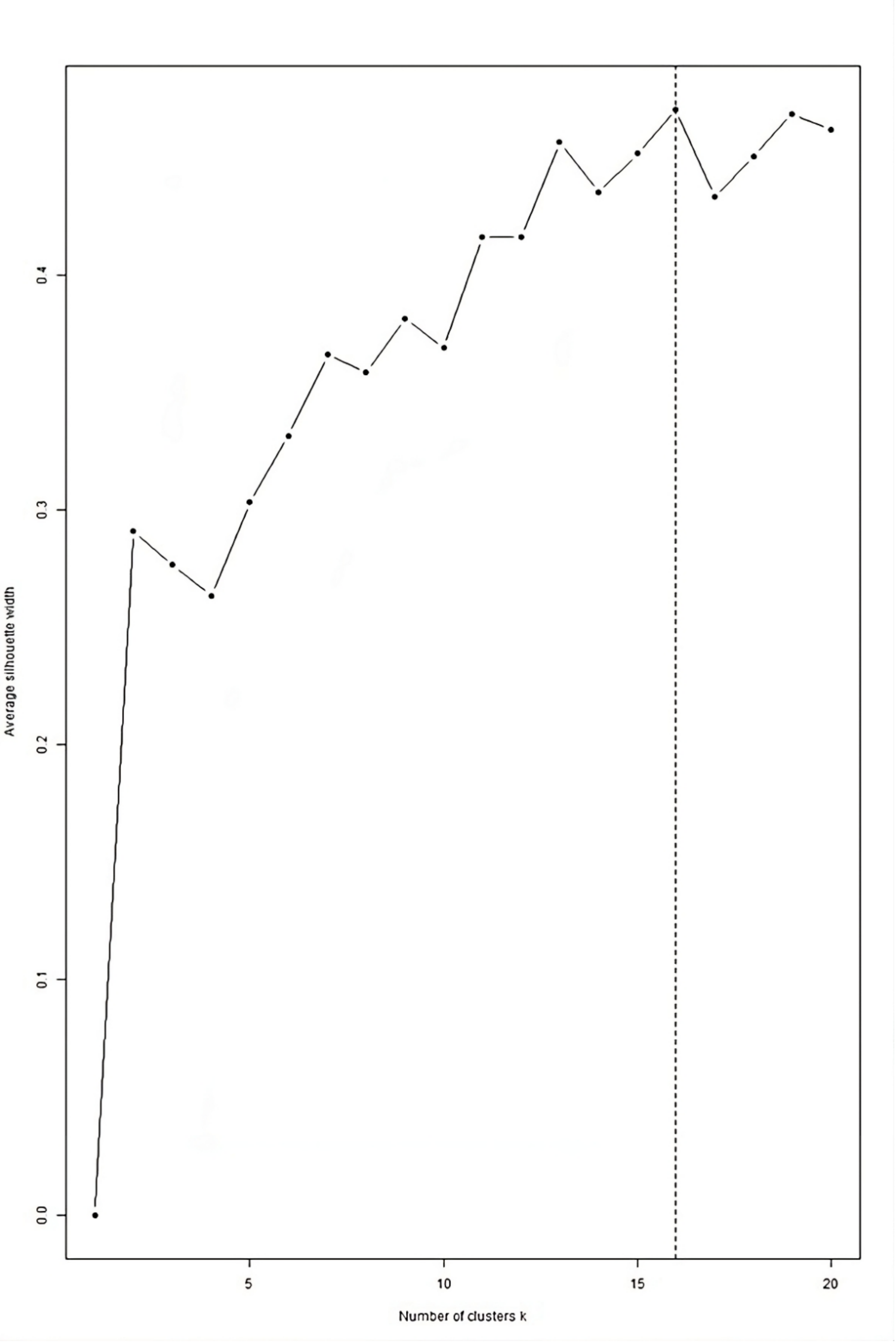

**Fig. S1** Average silhouette width (AWS) plot for hierarchical clustering of all samples. The X-axis represents the number of clusters, and the Y-axis is the average silhouette width. Cluster number $k = 16$ had the highest verage silhouette width. The closer the AWS value to 1, the more well-separated and distinct clusters are, suggesting that the data points within each cluster are similar to each other and dissimilar to points in other clusters

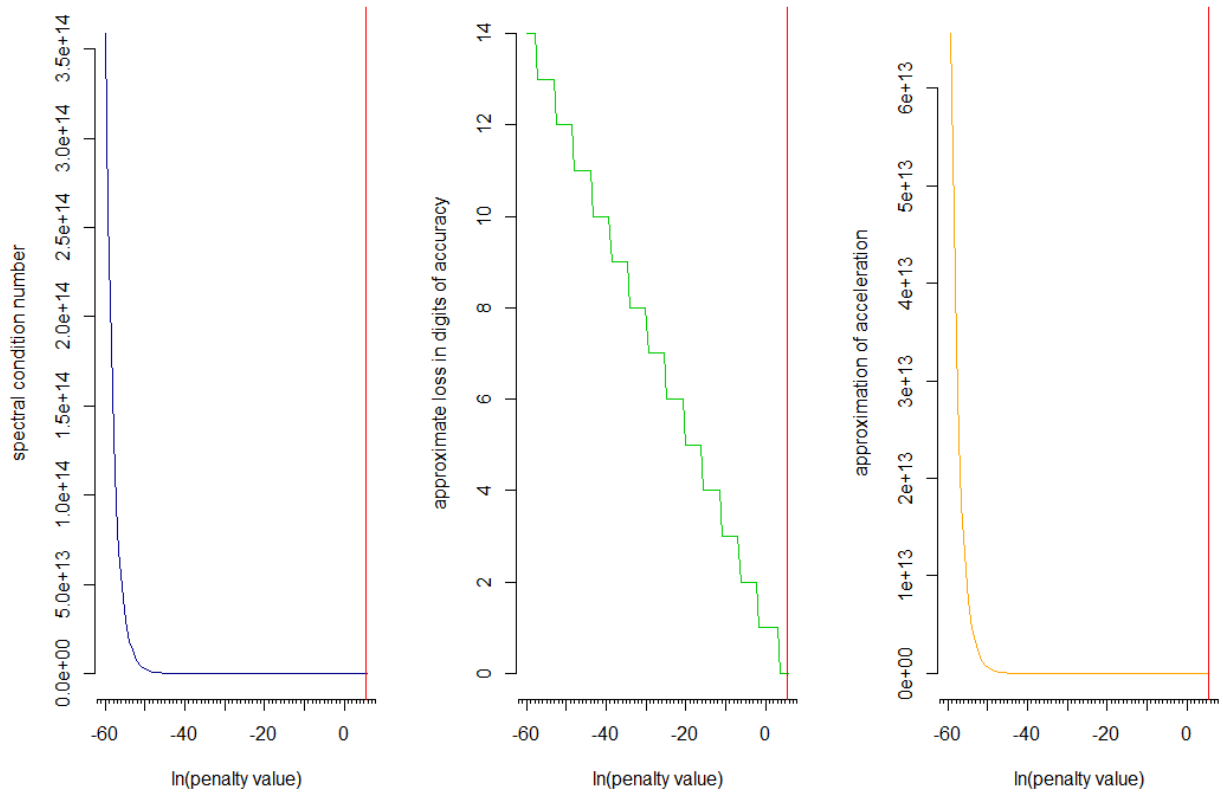

**Fig. S2** A condition number plot with interpretational aids for class Liver 30dpf. Left-handed panel: the condition number plot. Middle panel: the approximate loss in digits of accuracy (for the operation of inversion) [1] [2]. Right-handed panel: an approximation to the second-order derivative of the curvature in the basic plot [1] [2]. The vertical red line corresponds to the previously computed optimal ridge penalty for class Liver 30dpf. The condition number of a matrix measures how sensitive the matrix is to changes in its input – the numerically stable matrix is favourable, indicating higher accuracy and robustness. As seen in the left panel of the figure, the optimal penalty parameter exceeded the minimal value of the penalty parameter; the condition number remains relatively low and stable across the variations of $ln(penalty\ value)$ before reaching the estimated optimal penalty. An analogous pattern was present in pectral condition number plots of all class-specific ridges

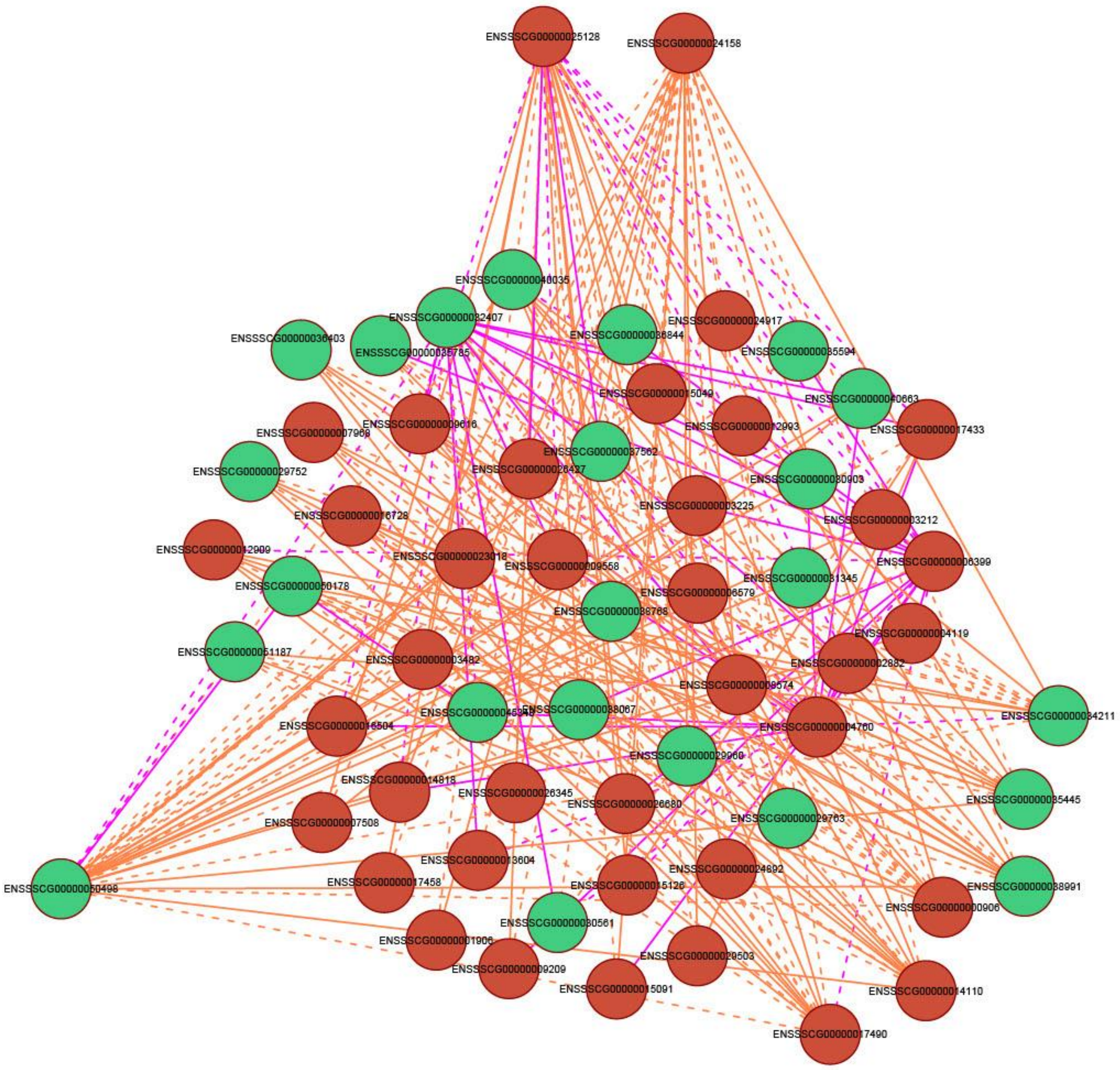

**Fig. S3** Differential network between Lung stage 70dpf and Lung stage NB. Red nodes indicate up-regulated genes and green down-regulated genes. Edges unique to class Lung 70dpf are visualized in pink, while edges unique to class Lung NB are in orange. Solid edges correspond to positive partial correlations, whereas dashed edges indicate negatively weighted partial correlations

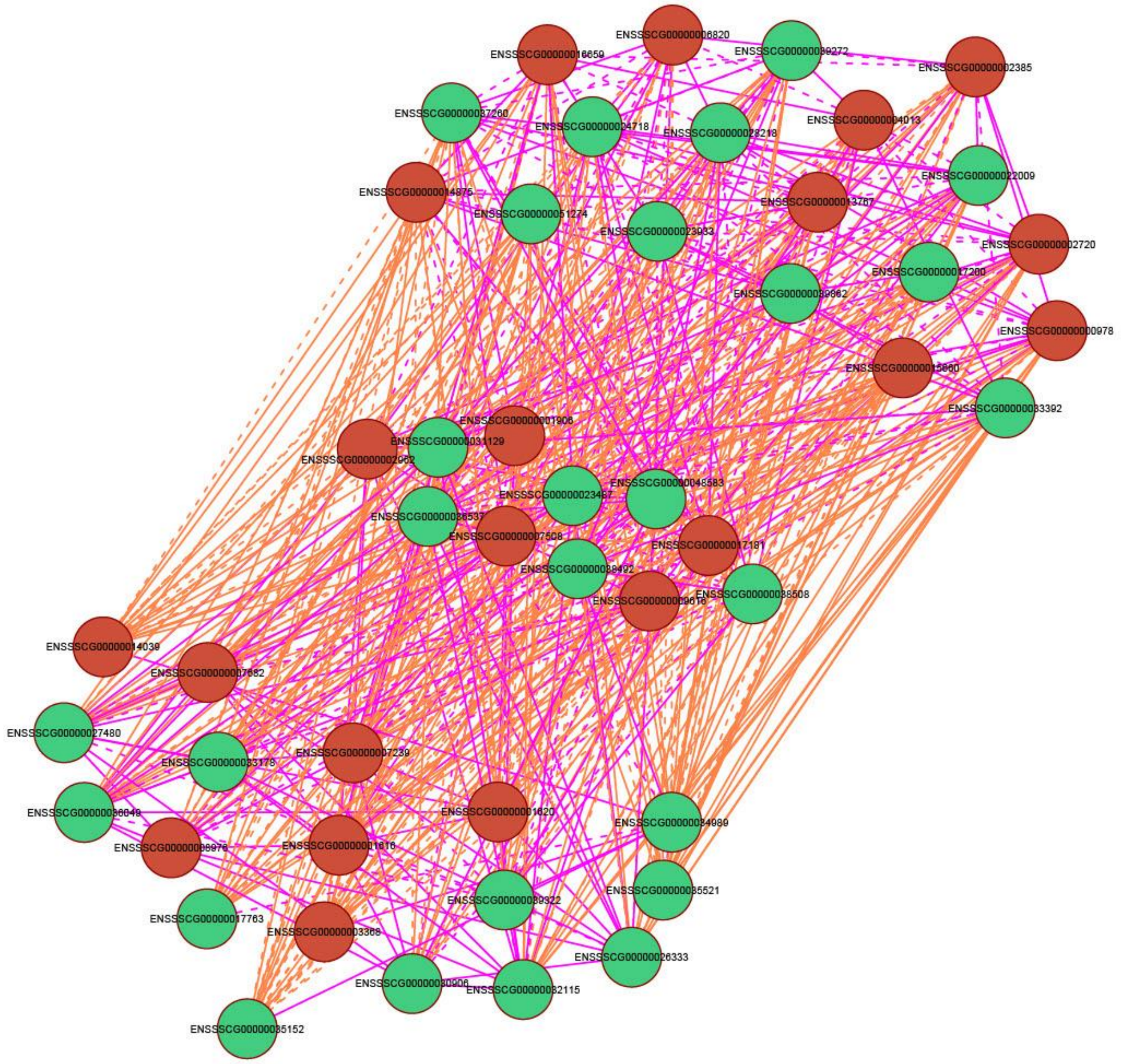

**Fig. S4** Differential network between Kidney stage 30dpf and Kidney stage 70dpf. Red nodes indicate up-regulated genes and green down-regulated genes. Edges unique to class Kidney 30dpf are visualized in pink, while edges unique to class Kidney 70dpf are in orange. Solid edges correspond to positive partial correlations, whereas dashed edges indicate negatively weighted partial correlations

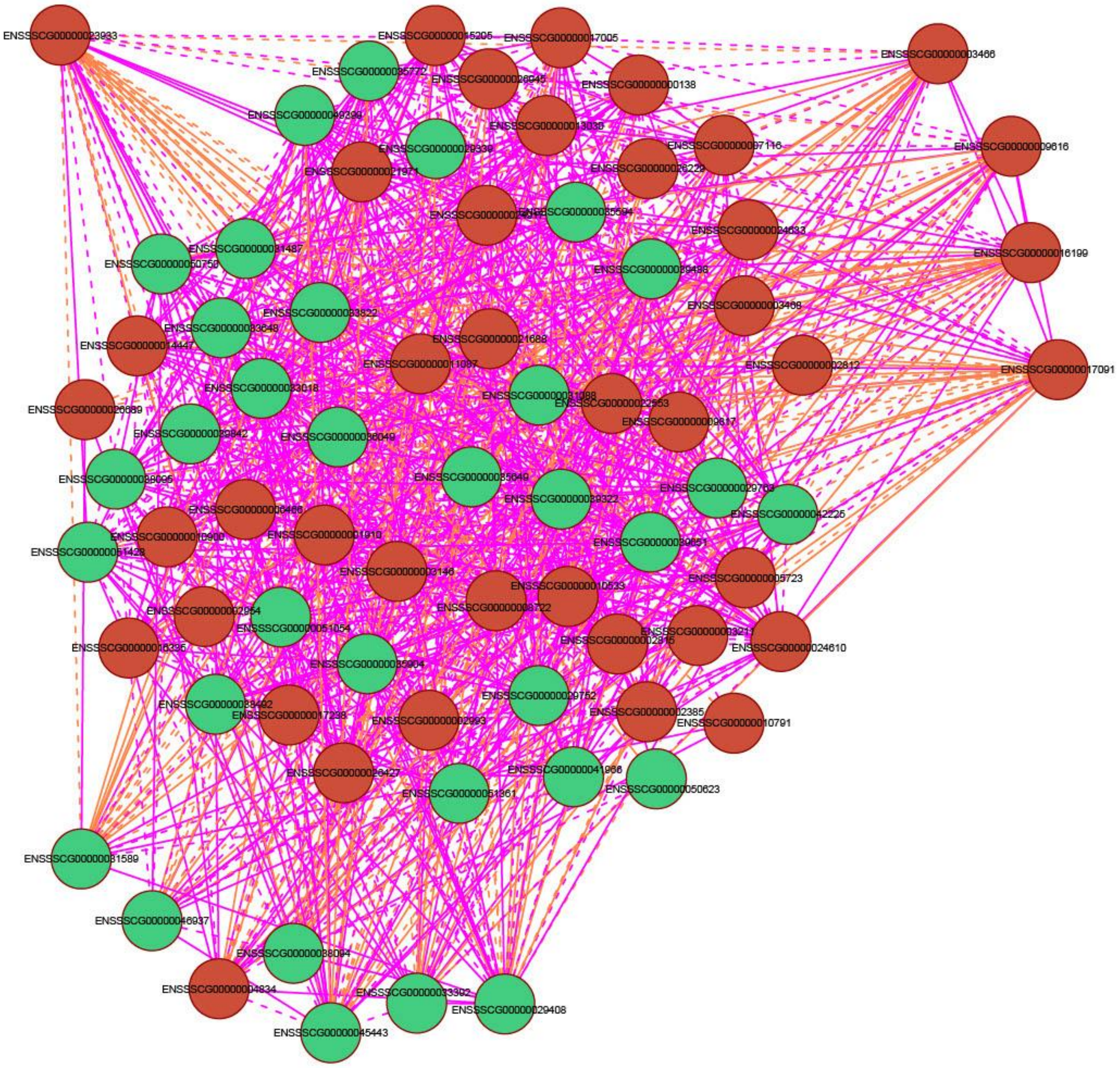

**Fig. S5** Differential network between Skin stage 70dpf and Skin stage NB. Red nodes indicate up-regulated genes and green down-regulated genes. Edges unique to class Skin 70dpf are visualized in pink, while edges unique to class Skin NB are in orange. Solid edges correspond to positive partial correlations, whereas dashed edges indicate negatively weighted partial correlations

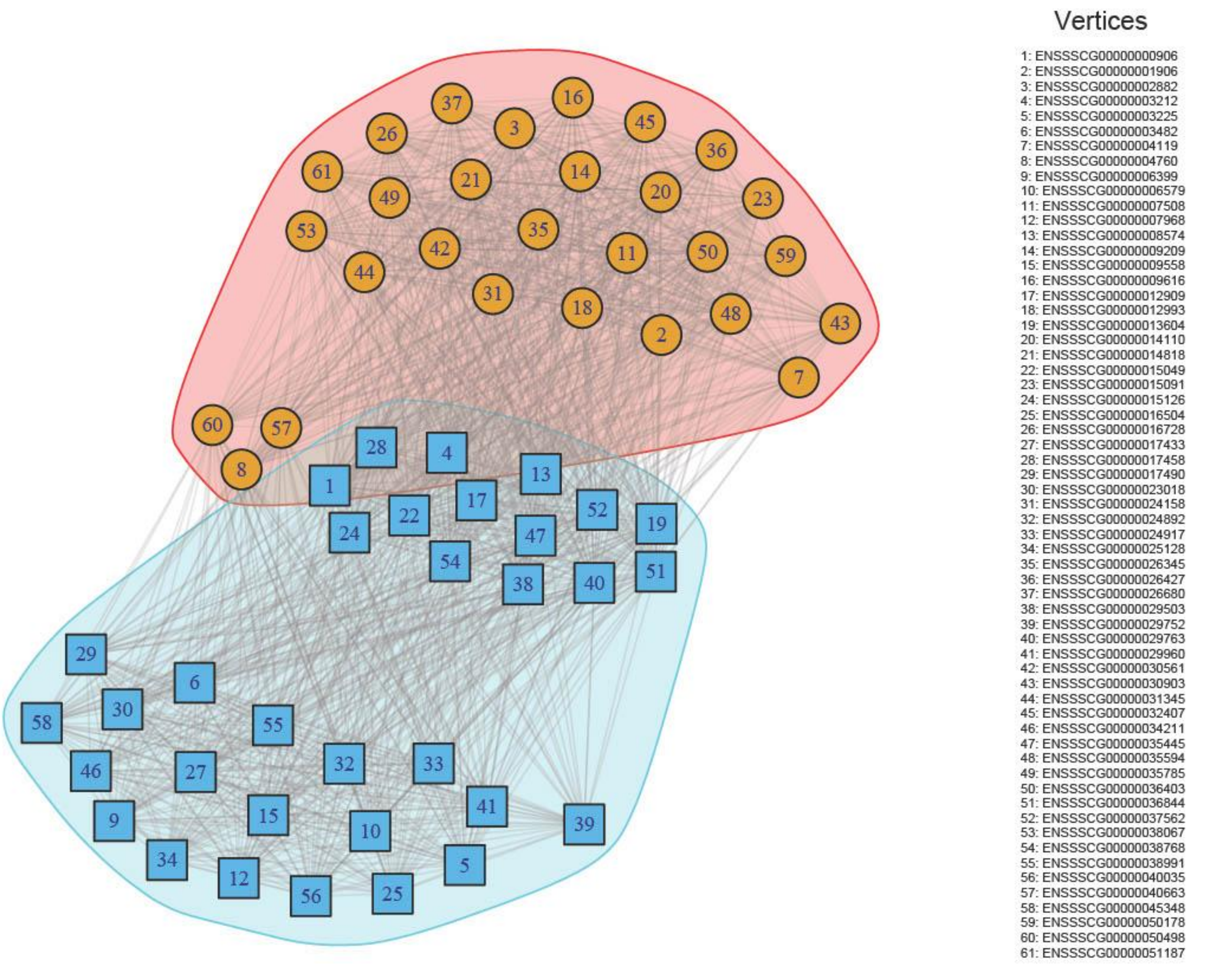

**Fig. S6** Communities detected for Liver 70dpf with the *cluster_fluid_communities()* function from the igraph R package. Each community is assigned a different colour. The legend on the right provides the encoding of Ensembl gene IDs in the network

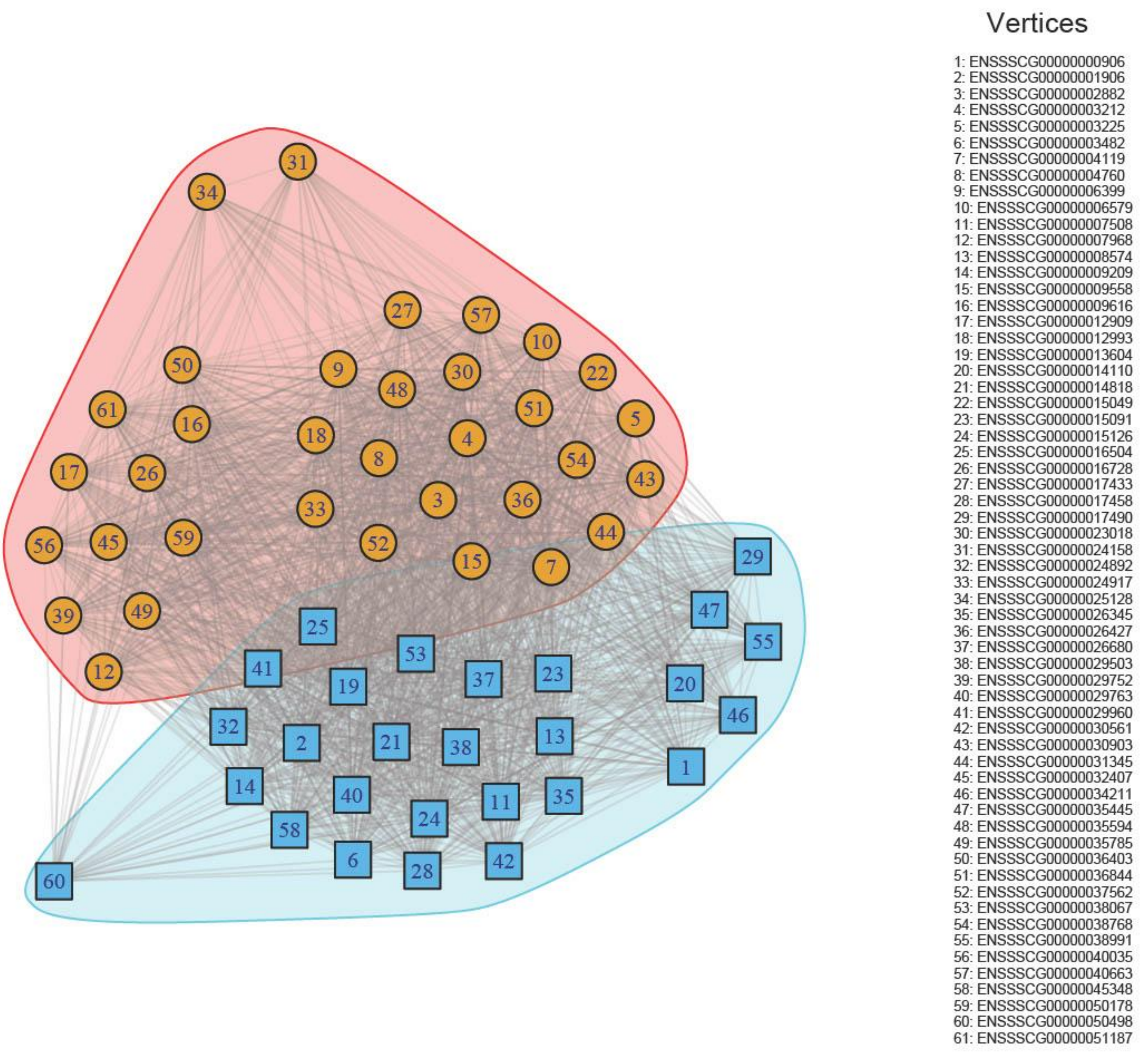

**Fig. S7** Communities detected for Lung 70dpf with the *cluster_fluid_communities()* function from the igraph R package. Each community is assigned a different colour. The legend on the right provides the encoding of Ensembl gene IDs in the network

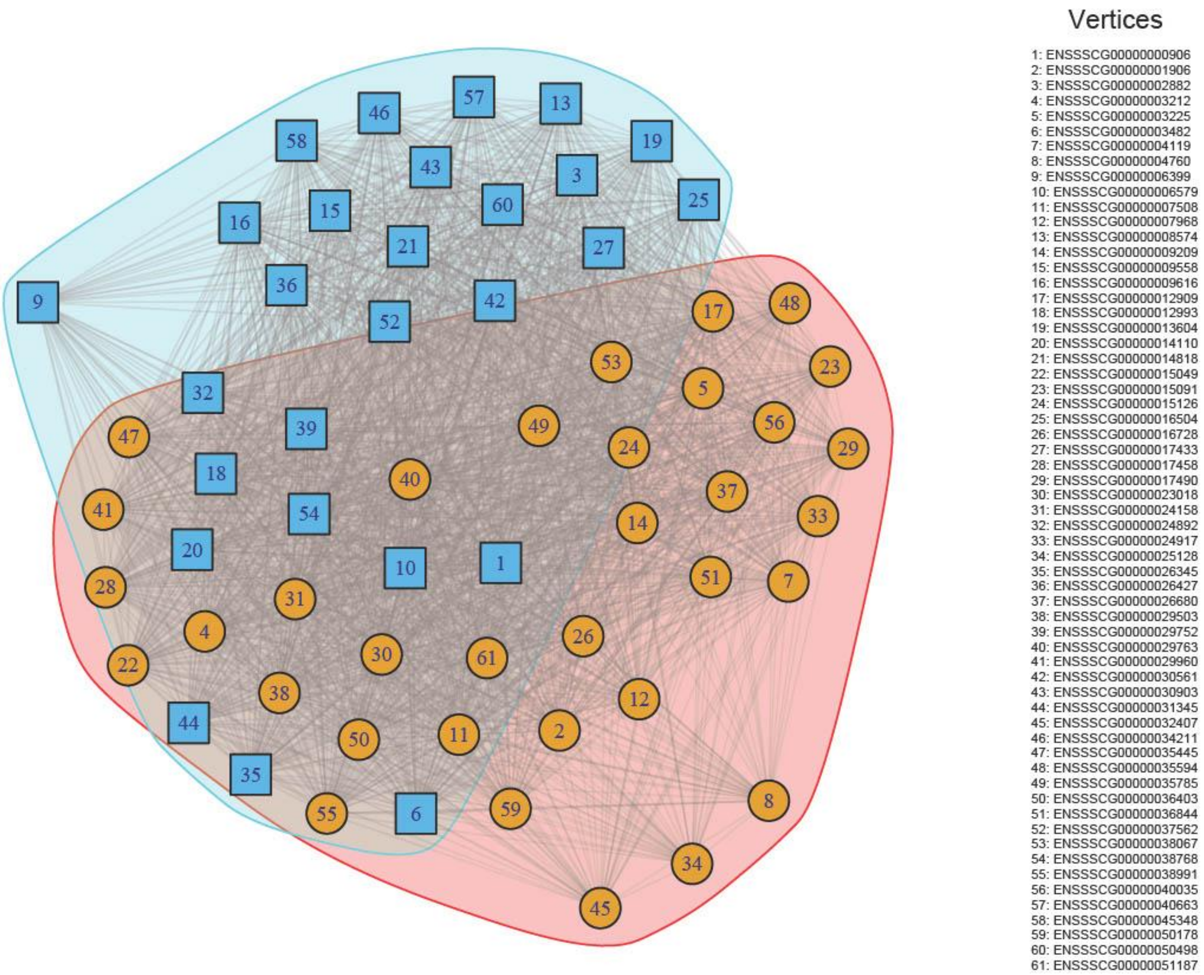

**Fig. S8** Communities detected for Lung NB with the *cluster_fluid_communities()* function from the igraph R package. Each community is assigned a different colour. The legend on the right provides the encoding of Ensembl gene IDs in the network

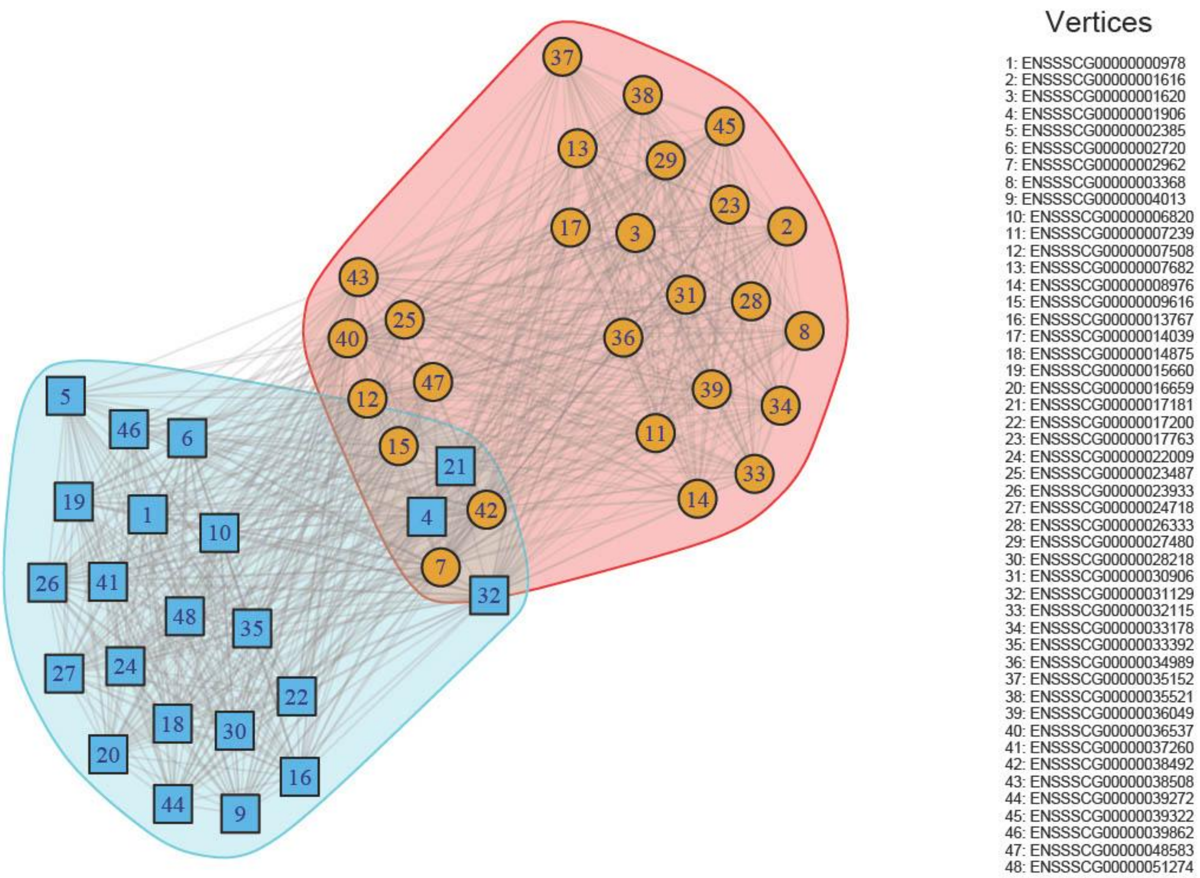

**Fig. S9** Communities detected for Kidney 30dpf with the *cluster_fluid_communities()* function from the igraph R package. Each community is assigned a different colour. The legend on the right provides the encoding of Ensembl gene IDs in the network

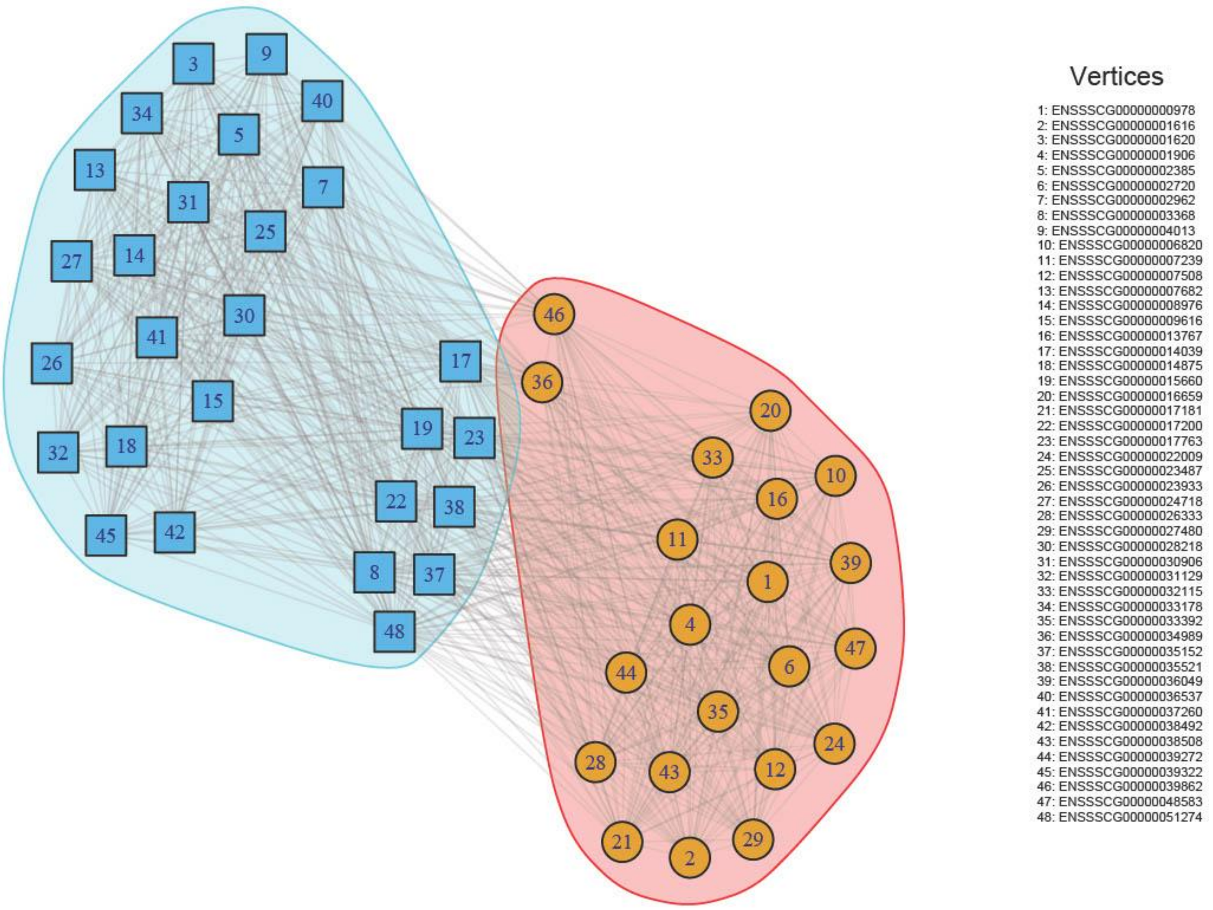

**Fig. S10** Communities detected for Kidney 70dpf with the *cluster_fluid_communities()* function from the igraph R package. Each community is assigned a different colour. The legend on the right provides the encoding of Ensembl gene IDs in the network

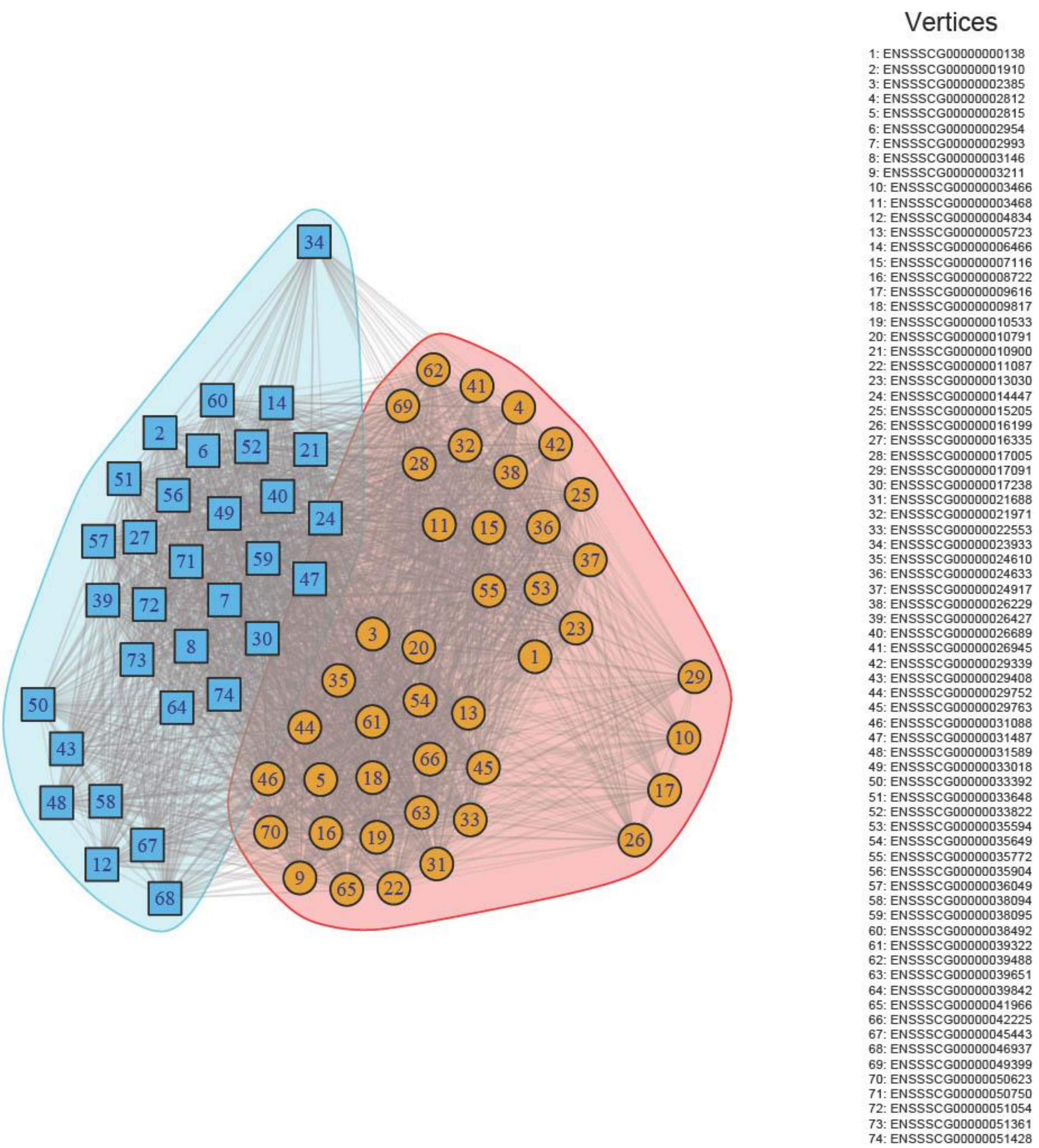

**Fig. S11** Communities detected for Skin 70dpf with the *cluster_fluid_communities()* function from the igraph R package. Each community is assigned a different colour. The legend on the right provides the encoding of Ensembl gene IDs in the network

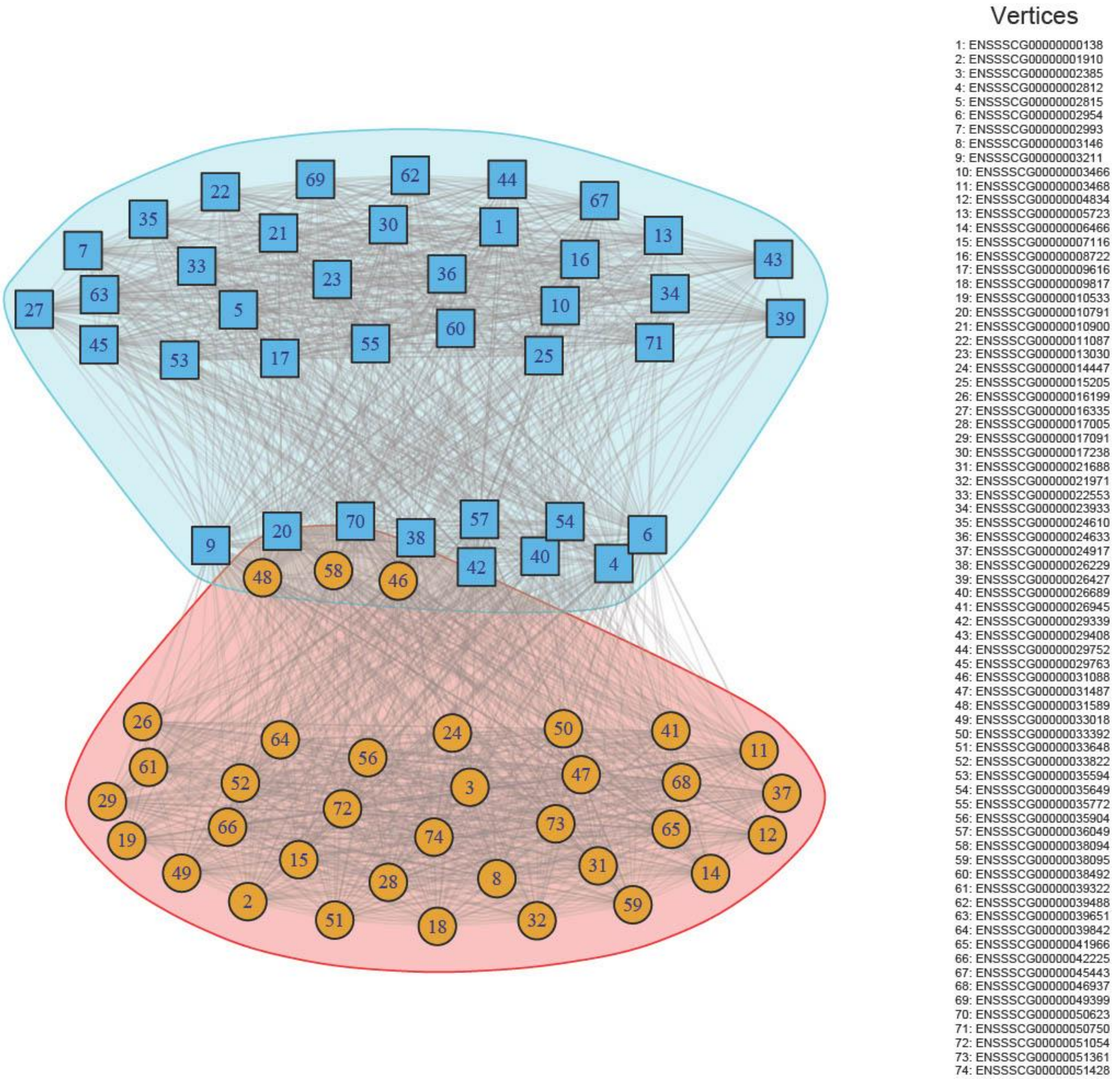

**Fig. S12** Communities detected for Skin NB with the *cluster_fluid_communities()* function from the igraph R package. Each community is assigned a different colour. The legend on the right provides the encoding of Ensembl gene IDs in the network

### TABLES

**Table S1:** An overview of all genes used for modeling endoderm-specific networks

| g# | initial alias | ortholog name | ortholog ensg | Description |
|---|---|---|---|---|
| 1 | ENSSSCG00000000906 | TMCC3 | ENSG00000057704 | transmembrane and coiled-coil domain family 3 |
| 2 | ENSSSCG00000001906 | CYP1A1 | ENSG00000140465 | cytochrome P450 family 1 subfamily A member 1 |
| 3 | ENSSSCG00000002882 | KRTDAP | ENSG00000188508 | keratinocyte differentiation associated protein |
| 4 | ENSSSCG00000003212 | NAPSA | ENSG00000131400 | napsin A aspartic peptidase |
| 5 | ENSSSCG00000003225 | CEACAM18 | ENSG00000213822 | CEA cell adhesion molecule 18 |
| 6 | ENSSSCG00000003482 | PADI3 | ENSG00000142619 | peptidyl arginine deiminase 3 |
| 7 | ENSSSCG00000004119 | GRM1 | ENSG00000152822 | glutamate metabotropic receptor 1 |
| 8 | ENSSSCG00000004760 | PPP1R14D | ENSG00000166143 | protein phosphatase 1 regulatory inhibitor subunit 14D |
| 9 | ENSSSCG00000006399 | VSIG8 | ENSG00000243284 | V-set and immunoglobulin domain containing 8 |
| 10 | ENSSSCG00000006579 | S100A3 | ENSG00000188015 | S100 calcium binding protein A3 |
| 11 | ENSSSCG00000007508 | ZBP1 | ENSG00000124256 | Z-DNA binding protein 1 |
| 12 | ENSSSCG00000007968 | RHBDF1 | ENSG00000007384 | rhomboid 5 homolog 1 |
| 13 | ENSSSCG00000008574 | KIF3C | ENSG00000084731 | kinesin family member 3C |
| 14 | ENSSSCG00000009209 | N/A | N/A | N/A |
| 15 | ENSSSCG00000009558 | F10 | ENSG00000126218 | coagulation factor X |
| 16 | ENSSSCG00000009616 | HR | ENSG00000168453 | HR lysine demethylase and nuclear receptor core- pressor |
| 17 | ENSSSCG00000012909 | PTPRCAP | ENSG00000213402 | protein tyrosine phosphatase receptor type C associ- ated protein |
| 18 | ENSSSCG00000012993 | SLC25A45 | ENSG00000162241 | solute carrier family 25 member 45 |
| 19 | ENSSSCG00000013604 | MYO1F | ENSG00000142347 | myosin IF |
| 20 | ENSSSCG00000014110 | DMGDH | ENSG00000132837 | dimethylglycine dehydrogenase |
| 21 | ENSSSCG00000014818 | STARD10 | ENSG00000214530 | StAR related lipid transfer domain containing 10 |
| 22 | ENSSSCG00000015049 | TMPRSS5 | ENSG00000166682 | transmembrane serine protease 5 |
| 23 | ENSSSCG00000015091 | MPZL2 | ENSG00000149573 | myelin protein zero like 2 |
| 24 | ENSSSCG00000015126 | TRIM29 | ENSG00000137699 | tripartite motif containing 29 |
| 25 | ENSSSCG00000016504 | TBXAS1 | ENSG00000059377 | thromboxane A synthase 1 |
| 26 | ENSSSCG00000016728 | IGFBP1 | ENSG00000146678 | insulin like growth factor binding protein 1 |
| 27 | ENSSSCG00000017433 | N/A | N/A | N/A |
| 28 | ENSSSCG00000017458 | KRT39 | ENSG00000196859 | keratin 39 |
| 29 | ENSSSCG00000017490 | GSDMA | ENSG00000167914 | gasdermin A |
| 30 | ENSSSCG00000023018 | MCCD1 | ENSG00000204511 | mitochondrial coiled-coil domain 1 |
| 31 | ENSSSCG00000024158 | ANO1 | ENSG00000131620 | anoctamin 1 |
| 32 | ENSSSCG00000024892 | NSG1 | ENSG00000168824 | neuronal vesicle trafficking associated 1 |
| 33 | ENSSSCG00000024917 | N/A | N/A | N/A |
| 34 | ENSSSCG00000025128 | CPN2 | ENSG00000178772 | carboxypeptidase N subunit 2 |
| 35 | ENSSSCG00000026345 | CLCNKB | ENSG00000184908 | chloride voltage-gated channel Kb |
| 35 | ENSSSCG00000026345 | CLCNKA | ENSG00000186510 | chloride voltage-gated channel Ka |
| 36 | ENSSSCG00000026427 | RORC | ENSG00000143365 | RAR related orphan receptor C |
| 37 | ENSSSCG00000026680 | KCND3 | ENSG00000171385 | potassium voltage-gated channel subfamily D mem- ber 3 |
| 38 | ENSSSCG00000029503 | F7 | ENSG00000057593 | coagulation factor VII |
| 39 | ENSSSCG00000029752 | C16orf54 | ENSG00000185905 | chromosome 16 open reading frame 54 |
| 40 | ENSSSCG00000029763 | IFI35 | ENSG00000068079 | interferon induced protein 35 |
| 41 | ENSSSCG00000029960 | LRRC4B | ENSG00000131409 | leucine rich repeat containing 4B |
| 42 | ENSSSCG00000030561 | LMTK3 | ENSG00000142235 | lemur tyrosine kinase 3 |
| 43 | ENSSSCG00000030903 | CDSN | ENSG00000204539 | corneodesmosin |

| | | | | |
|---|---|---|---|---|
| 44 | ENSSSCG00000031345 | INSYN2A | ENSG00000188916 | inhibitory synaptic factor 2A |
| 45 | ENSSSCG00000032407 | LRFN1 | ENSG00000128011 | leucine rich repeat and fibronectin type III domain containing 1 |
| 46 | ENSSSCG00000034211 | SPIB | ENSG00000269404 | Spi-B transcription factor |
| 47 | ENSSSCG00000035445 | SEZ6 | ENSG00000063015 | seizure related 6 homolog |
| 48 | ENSSSCG00000035594 | PRSS22 | ENSG00000005001 | serine protease 22 |
| 49 | ENSSSCG00000035785 | N/A | N/A | N/A |
| 50 | ENSSSCG00000036403 | FAM180A | ENSG00000189320 | family with sequence similarity 180 member A |
| 51 | ENSSSCG00000036844 | GPR171 | ENSG00000174946 | G protein-coupled receptor 171 |
| 52 | ENSSSCG00000037562 | SLC2A4RG | ENSG00000125520 | SLC2A4 regulator |
| 53 | ENSSSCG00000038067 | SMIM24 | ENSG00000095932 | small integral membrane protein 24 |
| 54 | ENSSSCG00000038768 | GPR37L1 | ENSG[2]00000170075 | G protein-coupled receptor 37 like 1 |
| 55 | ENSSSCG00000038991 | S100A13 | ENSG00000189171 | S100 calcium binding protein A13 |
| 56 | ENSSSCG00000040035 | MUC2 | ENSG00000198788 | mucin 2, oligomeric mucus/gel-forming |
| 57 | ENSSSCG00000040663 | HERPUD1 | ENSG00000051108 | homocysteine inducible ER protein with ubiquitin like domain 1 |
| 58 | ENSSSCG00000045348 | TMEM213 | ENSG00000214128 | transmembrane protein 213 |
| 59 | ENSSSCG00000050178 | N/A | N/A | N/A |
| 60 | ENSSSCG00000050498 | PURA | ENSG00000185129 | purine rich element binding protein A |
| 61 | ENSSSCG00000051187 | GARIN5B | ENSG00000180043 | golgi associated RAB2 interactor family member 5B |

*Note:* The initial Ensembl Sscrofa11.1 gene IDs were converted to human orthologs with the g:Profiler version e110_eg57_p18_4b54a898, database. The **initial_alias** column corresponds to Ensembl Sscrofa11.1 gene IDs. The **ortholog_name** column contains the human ortholog Entrez IDs, whereas **ortholog_ensg** contains Ensembl IDs of the human orthologs. The **description** column provides additional information on genes.

**Table S2:** An overview of all genes used for modeling mesoderm-specific networks

| g# | initial alias | ortholog name | ortholog ensg | Description |
|---|---|---|---|---|
| 1 | ENSSSCG00000000978 | MLC1 | ENSG00000100427 | modulator of VRAC current 1 |
| 2 | ENSSSCG00000001616 | TREML2 | ENSG00000112195 | triggering receptor expressed on myeloid cells like 2 |
| 3 | ENSSSCG00000001620 | MDFI | ENSG00000112559 | MyoD family inhibitor |
| 4 | ENSSSCG00000001906 | CYP1A1 | ENSG00000140465 | cytochrome P450 family 1 subfamily A member 1 |
| 5 | ENSSSCG00000002385 | TGFB3 | ENSG00000119699 | transforming growth factor beta 3 |
| 6 | ENSSSCG00000002720 | CLEC18B | ENSG00000140839 | C-type lectin domain family 18 member B |
| 6 | ENSSSCG00000002720 | CLEC18A | ENSG00000157322 | C-type lectin domain family 18 member A |
| 6 | ENSSSCG00000002720 | CLEC18C | ENSG00000157335 | C-type lectin domain family 18 member C |
| 7 | ENSSSCG00000002962 | MAP4K1 | ENSG00000104814 | mitogen-activated protein kinase kinase kinase kinase 1 |
| 8 | ENSSSCG00000003368 | RNF207 | ENSG00000158286 | ring finger protein 207 |
| 9 | ENSSSCG00000004013 | SMOC2 | ENSG00000112562 | SPARC related modular calcium binding 2 |
| 10 | ENSSSCG00000006820 | EPS8L3 | ENSG00000198758 | EPS8 like 3 |
| 11 | ENSSSCG00000007239 | CCM2L | ENSG00000101331 | CCM2 like scaffold protein |
| 12 | ENSSSCG00000007508 | ZBP1 | ENSG00000124256 | Z-DNA binding protein 1 |
| 13 | ENSSSCG00000007682 | SH2B2 | ENSG00000160999 | SH2B adaptor protein 2 |
| 14 | ENSSSCG00000008976 | ART3 | ENSG00000156219 | ADP-ribosyltransferase 3 (inactive) |
| 15 | ENSSSCG00000009616 | HR | ENSG00000168453 | HR lysine demethylase and nuclear receptor corepressor |
| 16 | ENSSSCG00000013767 | PALM3 | ENSG00000187867 | paralemmin 3 |
| 17 | ENSSSCG00000014039 | RGS14 | ENSG00000169220 | regulator of G protein signaling 14 |
| 18 | ENSSSCG00000014875 | CAPN5 | ENSG00000149260 | calpain 5 |
| 19 | ENSSSCG00000015660 | PFKFB2 | ENSG00000123836 | 6-phosphofructo-2-kinase/fructose-2,6-biphosphatase 2 |
| 20 | ENSSSCG00000016659 | MATCAP2 | ENSG00000164542 | microtubule associated tyrosine carboxypeptidase 2 |
| 21 | ENSSSCG00000017181 | CYGB | ENSG00000161544 | cytoglobin |
| 22 | ENSSSCG00000017200 | UNC13D | ENSG00000092929 | unc-13 homolog D |
| 23 | ENSSSCG00000017763 | SLC13A2 | ENSG00000007216 | solute carrier family 13 member 2 |

| | | | | |
|---|---|---|---|---|
| 24 | ENSSSCG00000022009 | DDC | ENSG00000132437 | dopa decarboxylase |
| 25 | ENSSSCG00000023487 | MSLNL | ENSG00000162006 | mesothelin like |
| 26 | ENSSSCG00000023933 | CRACR2B | ENSG00000177685 | calcium release activated channel regulator 2B |
| 27 | ENSSSCG00000024718 | RIBC2 | ENSG00000128408 | RIB43A domain with coiled-coils 2 |
| 28 | ENSSSCG00000026333 | GTSF1 | ENSG00000170627 | gametocyte specific factor 1 |
| 29 | ENSSSCG00000027480 | KLF10 | ENSG00000155090 | KLF transcription factor 10 |
| 30 | ENSSSCG00000028218 | CEACAM20 | ENSG00000273777 | CEA cell adhesion molecule 20 |
| 31 | ENSSSCG00000030906 | MPL | ENSG00000117400 | MPL proto-oncogene, thrombopoietin receptor |
| 32 | ENSSSCG00000031129 | SIRT5 | ENSG00000124523 | sirtuin 5 |
| 33 | ENSSSCG00000032115 | OSGIN1 | ENSG00000140961 | oxidative stress induced growth inhibitor 1 |
| 34 | ENSSSCG00000033178 | HAUS4 | ENSG00000092036 | HAUS augmin like complex subunit 4 |
| 35 | ENSSSCG00000033392 | SCML4 | ENSG00000146285 | Scm polycomb group protein like 4 |
| 36 | ENSSSCG00000034989 | LRRTM2 | ENSG00000146006 | leucine rich repeat transmembrane neuronal 2 |
| 37 | ENSSSCG00000035152 | TEF | ENSG00000167074 | TEF transcription factor, PAR bZIP family member |
| 38 | ENSSSCG00000035521 | KLHL38 | ENSG00000175946 | kelch like family member 38 |
| 39 | ENSSSCG00000036049 | PEG10 | ENSG00000242265 | paternally expressed 10 |
| 40 | ENSSSCG00000036537 | NFIC | ENSG00000141905 | nuclear factor I C |
| 41 | ENSSSCG00000037260 | N/A | N/A | N/A |
| 42 | ENSSSCG00000038492 | PHETA2 | ENSG00000177096 | PH domain containing endocytic trafficking adaptor 2 |
| 43 | ENSSSCG00000038508 | SPTBN2 | ENSG00000173898 | spectrin beta, non-erythrocytic 2 |
| 44 | ENSSSCG00000039272 | IP6K3 | ENSG00000161896 | inositol hexakisphosphate kinase 3 |
| 45 | ENSSSCG00000039322 | NRGN | ENSG00000154146 | neurogranin |
| 46 | ENSSSCG00000039862 | TRIB3 | ENSG00000101255 | tribbles pseudokinase 3 |
| 47 | ENSSSCG00000048583 | N/A | N/A | N/A |
| 48 | ENSSSCG00000051274 | N/A | N/A | N/A |

*Note:* The initial Ensembl Sscrofa11.1 gene IDs were converted to human orthologs with the g:Profiler version e110_eg57_p18_4b54a898, database. The **initial_alias** column corresponds to Ensembl Sscrofa11.1 gene IDs. The **ortholog_name** column contains the human ortholog Entrez IDs, whereas **ortholog_ensg** contains Ensembl IDs of the human orthologs. The **description** column provides additional information on genes.

**Table S3:** An overview of all genes used for modeling ectoderm-specific networks

| g# | initial alias | ortholog name | ortholog ensg | Description |
|---|---|---|---|---|
| 1 | ENSSSCG00000000138 | PVALB | ENSG00000100362 | parvalbumin |
| 2 | ENSSSCG00000001910 | ISLR | ENSG00000129009 | immunoglobulin superfamily containing leucine rich repeat |
| 3 | ENSSSCG00000002385 | TGFB3 | ENSG00000119699 | transforming growth factor beta 3 |
| 4 | ENSSSCG00000002812 | KIFC3 | ENSG00000140859 | kinesin family member C3 |
| 5 | ENSSSCG00000002815 | ADGRG1 | ENSG00000205336 | adhesion G protein-coupled receptor G1 |
| 6 | ENSSSCG00000002954 | SPINT2 | ENSG00000167642 | serine peptidase inhibitor, Kunitz type 2 |
| 7 | ENSSSCG00000002993 | CYP2A13 | ENSG00000197838 | cytochrome P450 family 2 subfamily A member 13 |
| 8 | ENSSSCG00000003146 | NTN5 | ENSG00000142233 | netrin 5 |
| 9 | ENSSSCG00000003211 | NR1H2 | ENSG00000131408 | nuclear receptor subfamily 1 group H member 2 |
| 10 | ENSSSCG00000003466 | TMEM82 | ENSG00000162460 | transmembrane protein 82 |
| 11 | ENSSSCG00000003468 | PLEKHM2 | ENSG00000116786 | pleckstrin homology and RUN domain containing M2 |
| 12 | ENSSSCG00000004834 | NDN | ENSG00000182636 | necdin, MAGE family member |
| 13 | ENSSSCG00000005723 | NTNG2 | ENSG00000196358 | netrin G2 |
| 14 | ENSSSCG00000006466 | SH2D2A | ENSG00000027869 | SH2 domain containing 2A |
| 15 | ENSSSCG00000007116 | CD93 | ENSG00000125810 | CD93 molecule |
| 16 | ENSSSCG00000008722 | SH3TC1 | ENSG00000125089 | SH3 domain and tetratricopeptide repeats 1 |
| 17 | ENSSSCG00000009616 | HR | ENSG00000168453 | HR lysine demethylase and nuclear receptor corepressor |
| 18 | ENSSSCG00000009817 | P2RX7 | ENSG00000089041 | purinergic receptor P2X 7 |
| 19 | ENSSSCG00000010533 | PYROXD2 | ENSG00000119943 | pyridine nucleotide-disulphide |

| | | | | |
|---|---|---|---|---|
| | | | | oxidoreductase domain 2 |
| 20 | ENSSSCG00000010791 | FUOM | ENSG00000148803 | fucose mutarotase |
| 21 | ENSSSCG00000010900 | DENND1B | ENSG00000213047 | DENN domain containing 1B |
| 22 | ENSSSCG00000011087 | SKIDA1 | ENSG00000180592 | SKI/DACH domain containing 1 |
| 23 | ENSSSCG00000013030 | PRDX5 | ENSG00000126432 | peroxiredoxin 5 |
| 24 | ENSSSCG00000014447 | SLC6A7 | ENSG00000011083 | solute carrier family 6 member 7 |
| 25 | ENSSSCG00000015205 | HEPACAM | ENSG00000165478 | hepatic and glial cell adhesion molecule |
| 26 | ENSSSCG00000016199 | CYP27A1 | ENSG00000135929 | cytochrome P450 family 27 subfamily A member 1 |
| 27 | ENSSSCG00000016335 | ESPNL | ENSG00000144488 | espin like |
| 28 | ENSSSCG00000017005 | KCNMB1 | ENSG00000145936 | potassium calcium-activated channel subfamily M regulatory beta subunit 1 |
| 29 | ENSSSCG00000017091 | TNIP1 | ENSG00000145901 | TNFAIP3 interacting protein 1 |
| 30 | ENSSSCG00000017238 | TTYH2 | ENSG00000141540 | tweety family member 2 |
| 31 | ENSSSCG00000021688 | ZDHHC13 | ENSG00000177054 | zinc finger DHHC-type palmitoyltransferase 13 |
| 32 | ENSSSCG00000021971 | DPEP1 | ENSG00000015413 | dipeptidase 1 |
| 33 | ENSSSCG00000022553 | TNRC18 | ENSG00000182095 | trinucleotide repeat containing 18 |
| 34 | ENSSSCG00000023933 | CRACR2B | ENSG00000177685 | calcium release activated channel regulator 2B |
| 35 | ENSSSCG00000024610 | KRT4 | ENSG00000170477 | keratin 4 |
| 36 | ENSSSCG00000024633 | JAKMIP3 | ENSG00000188385 | Janus kinase and microtubule interacting protein 3 |
| 37 | ENSSSCG00000024917 | N/A | N/A | N/A |
| 38 | ENSSSCG00000026229 | PCBD2 | ENSG00000132570 | pterin-4 alpha-carbinolamine dehydratase 2 |
| 39 | ENSSSCG00000026427 | RORC | ENSG00000143365 | RAR related orphan receptor C |
| 40 | ENSSSCG00000026689 | N/A | N/A | N/A |
| 41 | ENSSSCG00000026945 | LRRC2 | ENSG00000163827 | leucine rich repeat containing 2 |
| 42 | ENSSSCG00000029339 | SPATA2L | ENSG00000158792 | spermatogenesis associated 2 like |
| 43 | ENSSSCG00000029408 | ZFP30 | ENSG00000120784 | ZFP30 zinc finger protein |
| 44 | ENSSSCG00000029752 | C16orf54 | ENSG00000185905 | chromosome 16 open reading frame 54 |
| 45 | ENSSSCG00000029763 | IFI35 | ENSG00000068079 | interferon induced protein 35 |
| 46 | ENSSSCG00000031088 | N/A | N/A | N/A |
| 47 | ENSSSCG00000031487 | LSP1 | ENSG00000130592 | lymphocyte specific protein 1 |
| 48 | ENSSSCG00000031589 | ZNF37A | ENSG00000075407 | zinc finger protein 37A |
| 49 | ENSSSCG00000033018 | TM4SF1 | ENSG00000169908 | transmembrane 4 L six family member 1 |
| 50 | ENSSSCG00000033392 | SCML4 | ENSG00000146285 | Scm polycomb group protein like 4 |
| 51 | ENSSSCG00000033648 | N/A | N/A | N/A |
| 52 | ENSSSCG00000033822 | THRSP | ENSG00000151365 | thyroid hormone responsive |
| 53 | ENSSSCG00000035594 | PRSS22 | ENSG00000005001 | serine protease 22 |
| 54 | ENSSSCG00000035649 | N/A | N/A | N/A |
| 55 | ENSSSCG00000035772 | CDH5 | ENSG00000179776 | cadherin 5 |
| 56 | ENSSSCG00000035904 | RPL7A | ENSG00000148303 | ribosomal protein L7a |
| 57 | ENSSSCG00000036049 | PEG10 | ENSG00000242265 | paternally expressed 10 |
| 58 | ENSSSCG00000038094 | GPX7 | ENSG00000116157 | glutathione peroxidase 7 |
| 59 | ENSSSCG00000038095 | N/A | N/A | N/A |
| 60 | ENSSSCG00000038492 | PHETA2 | ENSG00000177096 | PH domain containing endocytic trafficking adaptor 2 |
| 61 | ENSSSCG00000039322 | NRGN | ENSG00000154146 | neurogranin |
| 62 | ENSSSCG00000039488 | SPON2 | ENSG00000159674 | spondin 2 |
| 63 | ENSSSCG00000039651 | SLC2A5 | ENSG00000142583 | solute carrier family 2 member 5 |
| 64 | ENSSSCG00000039842 | ZSCAN25 | ENSG00000197037 | zinc finger and SCAN domain containing 25 |
| 65 | ENSSSCG00000041966 | N/A | N/A | N/A |
| 66 | ENSSSCG00000042225 | N/A | N/A | N/A |
| 67 | ENSSSCG00000045443 | N/A | N/A | N/A |
| 68 | ENSSSCG00000046937 | N/A | N/A | N/A |

| | | | | |
|---|---|---|---|---|
| 69 | ENSSSCG00000049399 | N/A | N/A | N/A |
| 70 | ENSSSCG00000050623 | N/A | N/A | N/A |
| 71 | ENSSSCG00000050750 | N/A | N/A | N/A |
| 72 | ENSSSCG00000051054 | N/A | N/A | N/A |
| 73 | ENSSSCG00000051361 | N/A | N/A | N/A |
| 74 | ENSSSCG00000051428 | N/A | N/A | N/A |

*Note:* The initial Ensembl Sscrofa11.1 gene IDs were converted to human orthologs with the g:Profiler version e110_eg57_p18_4b54a898, database. The **initial_alias** column corresponds to Ensembl Sscrofa11.1 gene IDs. The **ortholog_name** column contains the human ortholog Entrez IDs, whereas **ortholog_ensg** contains Ensembl IDs of the human orthologs. The **description** column provides additional information on genes.
